# Harmonization of multi-site diffusion tensor imaging data

**DOI:** 10.1101/116541

**Authors:** Jean-Philippe Fortin, Drew Parker, Birkan Tunç, Takanori Watanabe, Mark A. Elliott, Kosha Ruparel, David R. Roalf, Theodore D. Satterthwaite, Ruben C. Gur, Raquel E. Gur, Robert T. Schultz, Ragini Verma, Russell T. Shinohara

**Affiliations:** Department of Biostatistics and Epidemiology, Perelman School of Medicine, University of Pennsylvania; Center for Biomedical Image Computing and Analytics, Department of Radiology, Perelman School of Medicine, University of Pennsylvania; Department of Radiology, Perelman School of Medicine, University of Pennsylvania; Department of Psychiatry, Perelman School of Medicine, University of Pennsylvania; Center for Autism Research, The Children’s Hospital of Philadelphia

**Keywords:** DTI, Diffusion, Harmonization, Multi-Site, ComBat, Inter-scanner

## Abstract

Diffusion tensor imaging (DTI) is a well-established magnetic resonance imaging (MRI) technique used for studying microstructural changes in the white matter. As with many other imaging modalities, DTI images suffer from technical between-scanner variation that hinders comparisons of images across imaging sites, scanners and over time. Using fractional anisotropy (FA) and mean diffusivity (MD) maps of 205 healthy participants acquired on two different scanners, we show that the DTI measurements are highly site-specific, highlighting the need of correcting for site effects before performing downstream statistical analyses. We first show evidence that combining DTI data from multiple sites, without harmonization, is counter-productive and negatively impacts the inference. Then, we propose and compare several harmonization approaches for DTI data, and show that ComBat, a popular batch-effect correction tool used in genomics, performs best at modeling and removing the unwanted inter-site variability in FA and MD maps. Using age as a biological phenotype of interest, we show that ComBat both preserves biological variability and removes the unwanted variation introduced by site. Finally, we assess the different harmonization methods in the presence of different levels of confounding between site and age, in addition to test robustness to small sample size studies.

## 1 Introduction

Diffusion tensor imaging (DTI) is a well-established magnetic resonance imaging (MRI) technique for studying the white matter (WM) organization and tissue characteristics of the brain. Diffusion tensor imaging has been used extensively to study both brain development and pathology; see Alexander et al. [2007] for a review of DTI and several of its applications. In studies assessing white matter tissue characteristics, two commonly reported complementary scalar maps are the mean diffusivity (MD), which assesses the degree to which water diffuses at each location, and fractional anisotropy (FA), which measures the coherence of this diffusion in one particular direction. Together, MD and FA provide complementary description of white matter microstructure.

With the increasing number of publicly availably neuroimaging databases, a crucial goal is to combine large-scale imaging studies to increase the power of statistical analyses to test common biological hypothesis. For instance, for life-span studies, combining data across sites and age ranges is essential for obtaining the necessary number of participants of each age. The success of combining multi-site imaging data depends critically on the comparability of the images across sites. As with other imaging modalities, DTI images are subject to technical variability across scans, including heterogeneity in the imaging protocol, variations in the scanning parameters and differences in the scanner manufacturers [Zhu et al., 2009, 2011]. Among others, the reliability of FA and MD maps have been shown to be affected by angular and spatial resolution [Zhan et al., 2010, Alexander et al., 2001, Kim et al., 2006], the number of diffusion weighting directions [Giannelli et al., 2009], the number of gradient sampling orientations [Jones, 2004], the number of b-values [Correia et al., 2009], and the b-values themselves.

In the design of multi-site studies, defining a standardized DTI protocol is a first step towards reducing inter-scanner variability. However, even in the presence of a standardized protocol, systematic differences between scanner manufacturers, field strength and other scanner characteristics will systematically affect the DTI images and induce inter-scanner variation. Image-based meta analysis (IBMA) techniques, reviewed in Salimi-Khorshidi et al. [2009], are common methods for combining results from multi-site studies with the goal of testing a statistical hypothesis. IBMA methods circumvent the need of harmonizing images across sites by performing site-specific statistical analyses and combining results afterwards. Fisher’s p-value combining method and Stouffer’s z-transformation test, applied to z or t-maps, are two common IBMA techniques. Fixed-effect models based on (possibly) normalized images, and mixed-effect models to model the inter- and intra-site variability, are other common techniques for the analysis of multi-site data. Indeed, meta-analysis methods have shown great promise for studies with a large number of participants at each site. For instance, the ENIGMA-DTI working group has been successfully using and validating meta-analysis techniques on such multi-site DTI data [Jahanshad et al., 2013, Kochunov et al., 2014].

Meta-analysis techniques have several limitations, however. First, study-specific samples might not be sufficient to estimate the true biological variability in the population [Mirzaalian et al., 2016]. As described by De Wit et al. [2014], adjusting for variability at the participant level is problematic in meta-analyses, since only group-level demographic and clinical information is available. Another limitation is that for a multi-site study, computing site-specific summary statistics will be affected by unbalanced data. For instance, the calculation of a variance using unbalanced datasets is highly affected by the ratio cases/controls in the sample [Linn et al., 2016b]. Another limitation, for imaging studies with small sample sizes, the parameters of the z-score transformations cannot be robustly estimated, yielding suboptimal statistical inferences.

Mega-analyses, in which the imaging data are combined before performing statistical inferences, have the potential to increase power compared to meta-analyses [De Wit et al., 2014]. In addition, pooling imaging data across studies has the benefit of enriching the clinical picture of the sample by increasing the variability in symptom profiles [Turner, 2014] and demographic variables. This is particularly important for age-span studies. However, pooling data across studies may increase the heterogeneity of the imaging measurements by introducing undesirable variability caused by differences in scanner protocols. Harmonization of the pooled data is therefore necessary to ensure the success of mega-analyses. The DTI harmonization technique proposed in Mirzaalian et al. [2016] is a first step towards that direction. The method is based on rotation invariant spherical harmonics (RISH) and combines the unprocessed DTI images across scanners. Unfortunately, a major drawback of the method is that it requires DTI data to have similar acquisition parameters across sites, an assumption often infeasible in multi-site observational analyses.

In this work, we adapt and compare several statistical approaches for the harmonization of DTI studies that were previously developed for other data types: Functional normalization [Fortin et al., 2014], RAVEL [Fortin et al., 2016a], Surrogate variable analysis (SVA) [Leek and Storey, 2007] and ComBat [Johnson et al., 2007], a popular batch adjustment method developed for genomics data. We also include a simple method that globally rescales the data for each site using a z-score transformation map common to all features, which we refer to as “global scaling”. For the evaluation of the different harmonization techniques, we use DTI data acquired as a part of two large imaging studies ([Satterthwaite et al., 2014] and [Ghanbari et al., 2014]) with images acquired on different scanners, using different imaging protocols. The participants are teenagers, and were matched across studies for age, gender, ethnicity, and handedness.

We first analyze site-related differences in the FA and MD measurements, and show evidence of significant site effects that differ across the brain. This motivates the need for a harmonization technique that is sensitive to region-specific scanner effects. Then, we harmonize the data with several proposed harmonizations, and evaluate their performance using a comprehensive evaluation framework. We consider a harmonization method to be successful if: (1) it removes the unwanted variation induced by site and differences in imaging protocols and (2) it preserves between-subject biological variability. As it is pointless to remove the noise associated with site if we cannot concurrently maintain the biological variation, both conditions must be simultaneously tested on the same set of images.

Using several criteria for evaluating (1) and (2), we show that the ComBat is the most effective harmonization technique for DTI studies, and is a promising method for other imaging studies. By allowing location-specific site correction factors and using an empirical Bayes framework, ComBat improves the estimation of the parameters for imaging sites with a small number of participants. It performs the best at removing unwanted variation induced by site while preserving the biological variation associated with age.

Similar to the other proposed methods in this paper, ComBat does not make assumptions about the origin of the site effects, and therefore does not require the images to be acquired with similar protocols.

## 2 Methods

### 2.1 Data

We consider two DTI studies from two different scanners. To investigate the effect of scanner variations on the DTI measurements, we matched the participants for age, gender, ethnicity and handedness, resulting in 105 participants retained in each study for further analysis. The characteristics of each dataset are described below.

#### Dataset 1 (Site 1): PNC dataset

We selected a subset of the Philadelphia Neurodevelopmental Cohort (PNC) [Satterthwaite et al., 2014], and included 105 healthy participants from 8 to 19 years old. 83 of the participants were males (22 females), and 75 participants were white (30 non-white). The DTI data were acquired on a 3T Siemens TIM Trio whole-body scanner, using a 32 channel head coil and a twice-refocused spin-echo (TRSE) single-shot EPI sequence with the following parameters: TR=8100 ms and TE = 82 ms, b-value of 1000 s/mm^2^, 7 b = 0 images and 64 gradient directions. The images were acquired at 1.875 × 1.875 × 2 mm resolution.

#### Dataset 2 (Site 2): ASD dataset

The dataset contains 105 healthy participants from a study focusing on autism spectrum disorder (ASD) [Ghanbari et al., 2014]. 83 of the participants were males (22 females), and 79 participants were white (26 non-white). The age of the participants ranges from 8 to 18 years old. The DTI data were acquired on a Siemens 3T Verio scanner, using a 32 channel head coil and a single shot spin-echo planar sequence with the following parameters: TR=11,000 ms and TE = 76 ms, b-value of 1000 s/mm^2^, 1 b = 0 image and 30 gradient directions. The images were acquired at 2mm isotropic resolution.

For benchmarking the different harmonization procedures, we use two additional subsets of the PNC database, with participants who differ from Dataset 1:

#### Independent Dataset 1

The dataset contains 292 additional healthy participants from the PNC with an age range similar to Dataset 1.

#### Independent Dataset 2

The dataset contains 105 additional healthy participants from the PNC with a slightly older age distribution than that of Dataset 1.

### 2.2 Image processing

Quality control on diffusion weighted images was performed manually. Datasets were excluded for field of view issues or intensity artifacts in the DWI that compromised more than 10% of the weighted images, or compromised the b0 image. Any artifact-affected weighted images were removed from the DWI volume if the volume as a whole was not excluded. DWI data were denoised using a joint anisotropic LMMSE filter for Rician noise [Tristán-Vega and Aja-Fernández, 2010]. The bo was extracted and skull-stripped using FSL’s BET tool [Smith, 2002], and the DTI model was fit within the brain mask using an unweighted linear least-squares method. Subsequently, the FA and MD maps were calculated from the resultant tensor image using the python package dipy [Garyfallidis et al., 2014]. The FA and MD images were co-registered to the T1-w image using FSL’s flirt tool [Jenkinson and Smith, 2001, Jenkinson et al., 2002]. The FA and MD maps were then non-linearly registered to the Eve template using DRAMMS [Ou et al., 2011]. A 3-tissue class T1-w segmentation was performed using FSL’s FAST tool [Zhang et al., 2001] to obtain GM, WM and CSF labels.

### 2.3 Harmonization methods

We propose to use and adapt five statistical harmonization techniques for DTI data: global scaling, functional normalization [Fortin et al., 2014], RAVEL [Fortin et al., 2016a], Surrogate Variable Analysis (SVA)[Leek and Storey, 2007, 2008] and ComBat [Johnson et al., 2007]. We refer to the absence of harmonization as “raw” data. We now describe the five different methods below with their implementation to the current datasets. For brevity, the different methods are presented in the context of FA intensities, but are similar for MD intensities and other modalities. We use the notation *y_ijυ_* to denote the FA measure at site *i*, for scan *j* and voxel *υ*.

#### 2.3.1 Global scaling

The global scaling (*GS*) approach is a model that assumes that the effect of each site on the DTI intensities can be summarized into a pair of a global shift and scale parameters (*θ_i,location_, θ_i,scale_*). More specifically, taking the average intensity map across all sites as a virtual reference site, the location parameter *θ_i,location_* and the scale parameter *θ_i,scale_* for site *i* can be obtained by fitting the linear model 
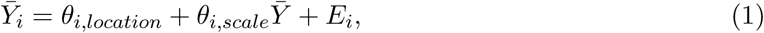
where 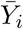 is the p×1 average vector of FA intensities for site *i*, 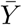 is the p × 1 global average vector of FA intensities across sites and *E_i_* is a vector of residuals assumed to have mean 0 and variance *σ ^2^*. Estimates 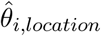 and 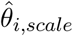 can be obtained by ordinary least squares (OLS). To remove the effect of site *i* on the data, we set the GS-harmonized FA intensity for voxel *v* and for scan *j* to be 
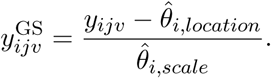

A more flexible model would be to fit a LOESS or LOWESS curve [Cleveland, 1979, 1981] at each site separately to allow for nonlinearity. This idea was previousy used in the so-called loess normalization [Bolstad et al., 2003].

#### 2.3.2 Functional normalization

We apply the functional normalization algorithm, described in Fortin et al. [2014] and later refined in Klein et al. [2015]. Unlike quantile normalization [Bolstad et al., 2003], which forces the histograms to be all the same across subjects, functional normalization only removes variation in the histograms that can be explained by a covariate. It has been successfully used to normalize cancer data and to normalize data from different genomic array technologies [Fortin et al., 2016b]. For multi-site DTI studies, we use site as a covariate. Briefly, the algorithm removes the site effect in the marginal distribution of the FA intensities by modeling the variation in the quantile functions as a function of site. After correction, the marginal densities of the FA intensities are more similar across sites.

#### 2.3.3 RAVEL

The RAVEL algorithm described in Fortin et al. [2016a] attempts to estimate a voxel-specific unwanted variation term by using a control region of the brain to estimate latent factors of unwanted variation common to all voxels. It is an extension of a previous intensity normalization, called White Stripe [Shinohara et al., 2014], developed to normalize white matter intensities in structural MRI. Similar to the control region used in Fortin et al. [2016a], we use voxels labelled as CSF as a control region. The FA values in CSF are expected to be 0, and fluctuations in the FA measurements in CSF are most likely technical in nature. In Figure A.1a), we show a strong correlation between average FA in the WM and average FA in the CSF. The FA values in the CSF can therefore act a surrogates for site effects in the WM.

Similar to RUV [Gagnon-Bartsch and Speed, 2012], we use singular value decomposition (SVD) to obtain *k* latent factors of unwanted variation, denoted w_1_,w_2_, …, w*_k_*, estimated from the CSF control voxels. Using cross-validation, we retain only the first latent factor for further analysis. At each voxel *v* in the WM, we fit the following linear regression model 
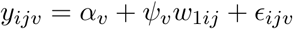
to obtain the voxel-specific RAVEL coefficients 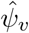 (shown in Figure A.1b in template space). We set the RAVEL-harmonized intensity to be 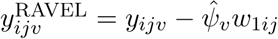. The results for MD maps are shown in Figure A.1c-d.

#### 2.3.4 SVA

The SVA algorithm estimates latent factors of unwanted variation, called surrogate variables, that are not associated with the biological covariates of interest. It is particularly useful when the site variable is not known, or when there exists residual unwanted variation after the removal of site effects. We used the reference implementation of SVA in the sva package [Leek et al., 2012], and surrogate variables were estimated using the iteratively re-weighted SVA algorithm [Leek and Storey, 2008]. We provided age and gender as covariates of interest to include in the regression models. The algorithm returns *s* surrogate variables z_1_, z_2_,*…*, z_*s*_, where *s* is estimated internally by the algorithm. Similar to RAVEL, we fit at each voxel *v* in the WM the following linear regression model x
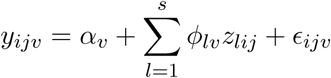
to obtain estimates 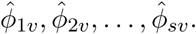 We set the SVA-harmonized intensity to be 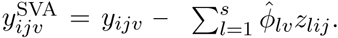

#### 2.3.5 ComBat

The ComBat model was introduced in the context of gene expression analysis by Johnson et al. [2007] as an improvement of location/scale models [Parmigiani et al., 2003] for studies with small sample size. Here, we reformulate the ComBat model in the context of DTI images. We assume that the data come from *m* imaging sites, containing each n_i_ scans within site *i* for *i* = 1,2,…, *m*, for voxel *v* = *1*,*2*,*…*, *p*. Let *y_ijv_* represent the FA measure for voxel *υ* for scan *j* for site *i*. After some standardization dicussed in Johnson et al. [2007], ComBat posits the following location and scale (L/S) adjustment model: 
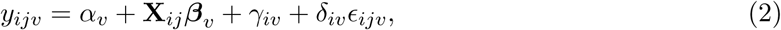
where *α_v_* is the overall FA measure for voxel *v*, X is a design matrix for the covariates of interest (e.g. gender, age), and (*β_v_* is the voxel-specific vector of regression coefficients corresponding to X. We further assume that the error terms *∈_ijv_* follow a normal distribution with mean zero and variance 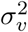. The terms *γ_iυ_* and *δ_iυ_* represent the additive and multiplicative site effects of site *i* for voxel *υ*, respectively. ComBat uses an empirical Bayes (EB) framework to estimate the parameters *γ_iυ_* and *δ_ig_* by pooling information across voxels. To do so, the site-effect parameters are assumed to have the parametric prior distributions: 
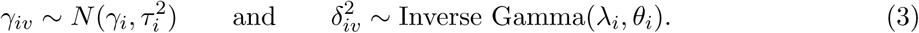

The hyperparameters 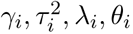 are estimated empirically from the data as described in Johnson et al. [2007]. In Figure 1, we present the distributions of the voxel-wise estimates *γ_iv_* and 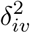 for each site (dotted lines) together with the estimated prior distributions (solid lines); the estimated prior distributions fit the data well. We note that the sva package also offers the option to posit non-parametric priors for more flexibility, at the cost of increasing computational time. As described in Johnson et al. [2007], the ComBat estimates 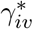 and 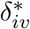 of the site effect parameters are computed using conditional posterior means, and are shown in Figure 1c) in template space.

**Figure 1.**
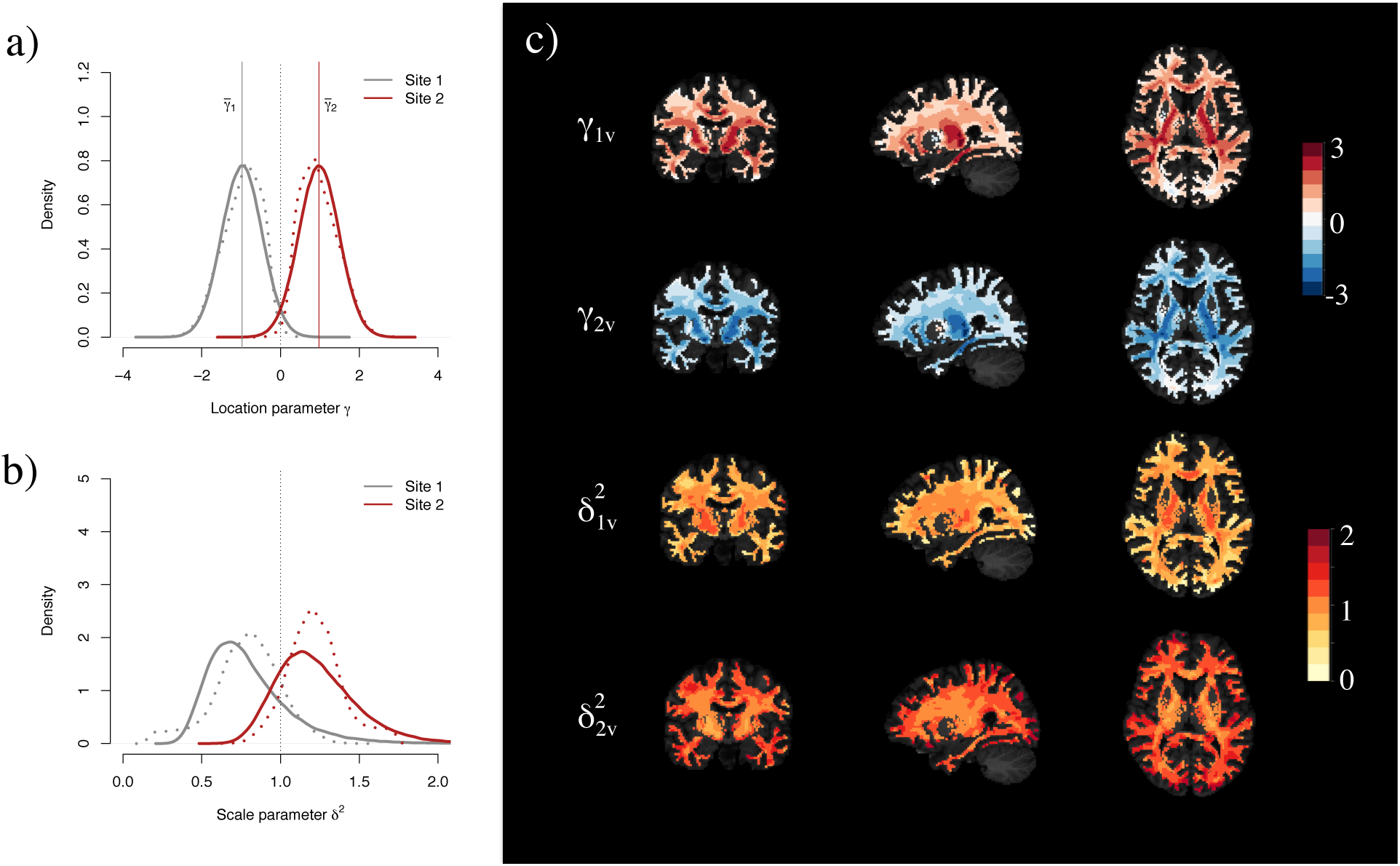
ComBat site effect parameters for FA. **(a)** The voxel-wise estimates of the location parameter *γ_iv_* for site 1 (dotted grey line) and site 2 (dotted red line) for the FA maps. The solid lines represent the prior distributions (normal distributions with mean 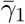 and 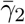 respectively) estimated in the ComBat procedure using empirical Bayes. **(b)** The voxel-wise estimates of the scale parameter *δ_iυ_* for site 1 (dotted grey line) and site 2 (dotted red line). The solid lines represent the EB-based prior distributions (inverse gamma distributions) estimated in the ComBat procedure. **(c)** Final EB-estimates for the site effects parameters for site 1 (first and third row) and site 2 (second and fourth row) in template space.

The final ComBat-harmonized FA values are defined as

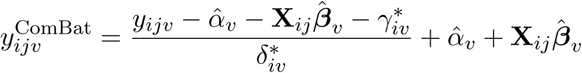

### 2.4 Evaluation framework

We consider a harmonization method to be successful if: (1) it removes the unwanted variation induced by site, scanner or differences in imaging protocols; (2) it preserves between-subject biological variability. Both conditions must be simultaneously tested on the same set of images; it is pointless to remove the noise associated with site if we cannot concurrently maintain the biological variation.

To evaluate (1), we calculate two-sample t-tests on the DTI intensities, testing for a difference between Site 1 and Site 2 measurements. We perform the analysis both at the voxel and ROI level. A harmonization technique that successfully removes site effect will result in non-significant tests, after possibly correcting for multiple comparisons. Criteria for (2) are harder to devise because of the absence of a gold-standard for the association of voxels or regions of the brain with a biological covariate. In Mirzaalian et al. [2016], the authors perform a multi-site harmonization on healthy participants, and show that the coefficient of variation (CoV) in FA at each site, measuring the intra-site variability, is comparable before and after harmonization. However, since the intra-site variability is a mixture of intra-site technical variability and intra-site biological variability, it does not directly follow that biological variability is preserved. Another approach to test whether or not (2) is satisfied is to create a synthetic experiment in which known biological effects, such as changes in FA associated with a particular disease, are added to a subset of the images. While potentially helpful, such simulations are rarely realistic; in real data, the structure of the noise is often complex and confounded with the signal component. Instead, we base our evaluation of (2) on the consistency, replicability and validity of voxels associated with biological variation, using age as the biological factor of interest. We further detail these criteria below.

#### 2.4.1 Consistency of the voxels associated with age

We use the consistency of the voxel-specific associations with age as a first primary criterion for the preservation of biological signal. Associations with age are measured using usual Wald t-statistics from linear regression. By consistency, we mean that the results returned by performing a statistical analysis at each site separately should be consistent with the results returned by performing a statistical analysis on the combined and harmonized data. The combined dataset is a mixture of both sites, and therefore the results should be correlated with the results obtained at each individual site. If it is not the case, this would indicate that combining datasets is counterproductive and adds noise to the analysis. We test the consistency of the voxels associated with age using a discovery-validation scheme.

In the discovery-validation scheme, we consider the harmonized dataset (Site 1 + Site2 + Harmonization) as a discovery cohort, and perform a mass-univariate analysis that estimates a t-statistic at each voxel testing for association with age. We denote the resulting vector of t-statistics by *t_dis_*. We then perform a mass-univariate analysis for the two sites separately, that we consider as two independent validation cohorts, and obtain two vectors of t-statistics *t_val1_* and *t_val2_*. in the case of a successful harmonization, the vector *t_dis_* should be more similar to both vectors *t_val1_* and *t_val2_*. While one could use the usual Pearson correlation coefficient to test for consistency, this has the drawback of considering all voxels equally (both signal and noise voxels), and therefore is not restrained to the voxels of interest. Because we wish to test for the consistency of the signal voxels only (voxels associated with age), we instead use concordance at the top (CAT) curves [Irizarry et al., 2005]. The CAT curves estimate the overlap between the top *k* t-statistics, which are the voxels most likely associated with age, for all possible values of *k*. *A CAT* curve closer to 1 indicates better overlap between the two lists of t-statistics. We calculate two CAT curves: CAT_1_, which measures the consistency between *t_dis_* and *t_val1_*, and *CAT*_2_, which measures the consistency between *t_dis_* and *t_val2_*. We consider the average curve 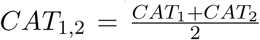 as a measure of consistency across studies. We summarize the discovery-validation scheme for consistency in Figure 2.

**Figure 2.**
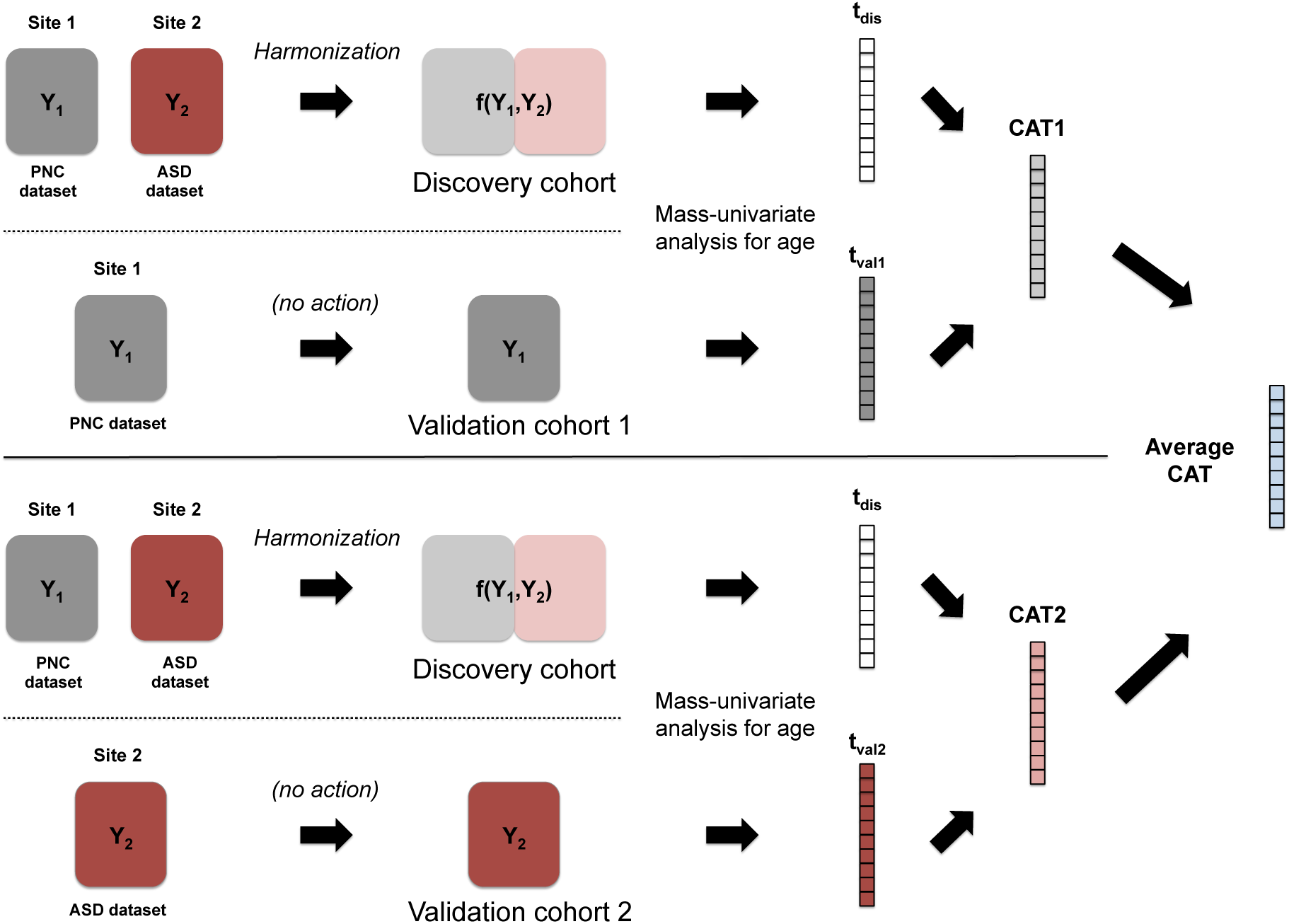
Discovery-validation scheme for the estimation of consistency. To estimate the performance of a harmonization procedure at improving the consistency of the voxels associated with age, we use the harmonized dataset as a discovery cohort, and both Site 1 and Site 2 separately as validation cohorts. For each cohort separately, we perform a mass-univariate analysis for age to obtain a t-statistic at each voxel. This yields one vector of t-statistics for the discovery cohort, *t_dis_*, and two vectors of t-statistics for the two validation cohorts (*t_val1_* for Site 1 and *t_val2_* for Site 2). We calculate the agreement between *t_dis_* and *t_val1_* using the concordance at the top (CAT) curve, described in the Methods section, and denote it by *CAT1*. Similarly, we calculate *CAT2* from *t_dis_* and *t_val2_*. A harmonization method that improves the consistency of the harmonized voxels with respect to the individual sites will improve the average CAT curve *(CAT1* + *CAT2*)*/*2.

#### 2.4.2 Replicability of the voxels associated with age

Another criterion for estimating the performance of image harmonization is to measure the replicability of the voxels associated with age. Replicability is defined as the chance that an independent experiment will produce a similar set of results [Leek and Peng, 2015], and is a strong indication that a set of results is biologically meaningful. We estimate the replicability of the voxels associated with age in a similar way as the consistency, presented above, but use independent discovery and validation cohorts. To achieve this, we consider the harmonized dataset (Site 1 + Site 2 + Harmonization) as a discovery cohort, and a independent dataset, with unrelated participants, as a validation cohort (Independent Dataset). Two independent datasets are considered in this paper and are described in the Data section. We then perform a mass-univariate analysis for the discovery and validation cohort separately, which yields two vectors of t-statistics, *t_dis_* and *t_val_*. We measure the overlap between *t_dis_* and *t_val_* using a CAT curve, which serves as a measure of replicability. We summarize the discovery-validation scheme for consistency in Figure 3.

**Figure 3.**
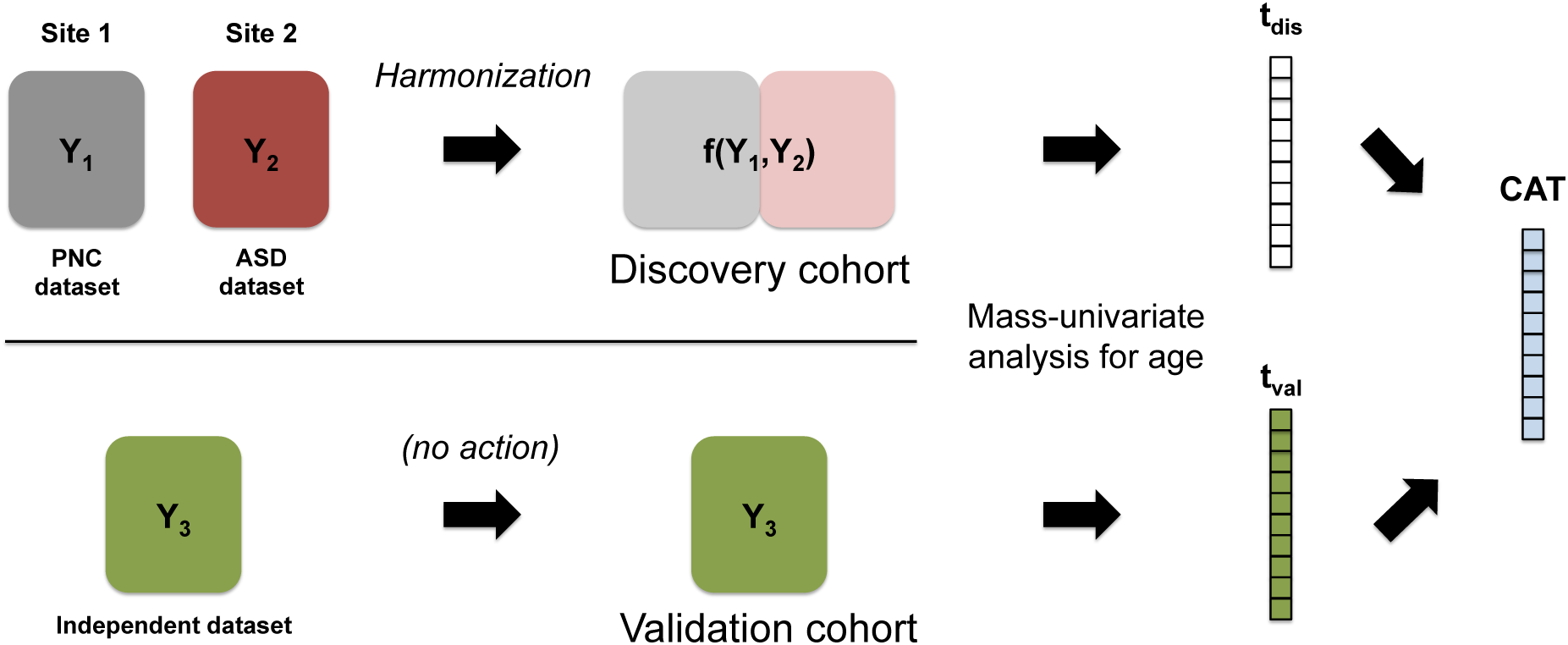
Discovery-validation scheme for the estimation of replicability. To estimate the performance of a harmonization procedure at improving the replicability of the voxels associated with age, we use the harmonized dataset as a discovery cohort, and an independent dataset (different participants) as a validation cohort. For each cohort separately, we perform a mass-univariate analysis for age to obtain a t-statistic at each voxel. This yields two vectors of t-statistics, *t_dis_* and *t_val_*, for the discovery and validation cohorts respectively. We calculate the agreement between *t_dis_* and *t_val_* using the concordance at the top (CAT) curve, described in the Methods section. A harmonization method that performs better will yield a vector *t_dis_* more similar to *t_val_*, that is a CAT curve closer to 1.

#### 2.4.3 Creation of silver-standards

To further evaluate the performance of the different harmonization methods, we create two sets of silver-standards: a silver-standard for voxels that are truly associated with age (signal silver-standard), and one for voxels not associated with age (null silver-standard).

##### Creation of a silver-standard for voxels associated with age

Many regions in the WM have been previously demonstrated to show an increase of FA values through adolescence, accompanied by decreasing of MD values [Tamnes et al., 2010, Bava et al., 2010, Lebel and Beaulieu, 2011]. Because some of the reported regions are specific to FA only, or specific to MD only, we estimate a reference set of voxels that substantially change with age for FA, and an additional set for substantial changes in the MD maps, for each site separately. Because our reference sets are estimated within site, they are free of site effects and should represent the best silver-standards for voxels associated with age: we refer to those sets as *signal silver-standards*. To estimate the signal silver-standard for FA (and similarly for MD), we use the following meta-analytic approach: for each site separately, at each voxel in the WM, we apply a linear regression model to obtain a t-statistic measuring the association of FA with age. For each site, we define the site-specific signal silver-standard to be the *k* = 5000 voxels with the highest t-statistics in magnitude. We define the signal silver-standard to be the intersection of the two site-specific signal silver standards. This ensures that the resulting voxels are not only voxels highly associated with age within a study, but are also replicated across the two sites.

For the FA values, this resulted in 2265 voxels for the signal silver-standard set. Among those voxels, 21.3% are located in the thalamic region, 17.1% are located in the anterior limb of the internal capsule (left and right) 14.8% in the posterior limb of the internal capsule (left and right), 10.8% in the midbrain, 9.7% in the cerebral peduncle and 4.9% in the globus pallidus. These results are highly consistent with the changes reported in literature for the same age group [Schmithorst et al., 2002, Barnea-Goraly et al., 2005, Ashtari et al., 2007, Bava et al., 2010, Giorgio et al., 2010]. For the MD values, we obtained a signal silver-standard set of 1932 voxels. 30.4% of these voxels are located in the superior corona radiata, 15.0% are located in the superior frontal lobe, 10.1% are located in the precentral region, 9.4% are located in the superior longitudinal fasciculus, 7.9% are located in the middle frontal region and 6.4% are located in the thalamic region, which is consistent with regions previously reported in the literature [Bava et al., 2010, Tamnes et al., 2010, Krogsrud et al., 2016].

##### Creation of a silver-standard for voxels not associated by age

In addition to a signal silver-standard for voxels associated with age, we created silver-standard sets for voxels unaffected by age, for both FA and MD maps, that we refer to as *null silver-standards*. Our approach is similar to that of signal silver-standards. For each site separately, at each voxel in the WM, we apply a linear regression model to obtain a t-statistic measuring the association of FA with age. For each site, we define the site-specific silver-standard for null voxels to be the *k* = 5000 voxels with the lowest t-statistics in magnitude (close to 0). We define the silver-standard to be the intersection of the two site-specific silver standards. This ensures that the resulting voxels are voxels with least association with age within a study, and are also replicated across the two sites.

For the FA values, we obtained a silver-standard set of 405 voxels. We note that this replication rate (8.1%) is much more lower than the replication rate for the signal silver-standard set (45.3%). This is not surprising; strong signal voxels are more likely to replicate than noise voxels. The top regions represented in the null silver-standard are the middle frontal lobe (8.6%), the middle occipital lobe (6.9%) and the precuneus region (5.4%). For the MD values, we obtained a null silver-standard set of 101 voxels. The top regions represented in the null silver-standard are the postcentral gyrus (5.5%) and the lingual gyrus (4.8%).

## 3 Results

The results are organized as follows. We first show evidence of substantial site effects in the FA and MD maps in Section 3.1, and then show how the different harmonization methods perform at removing those site effects in Section 3.2. In Section 3.3, we discuss the biological variability at each site separately, before and after harmonization and show how site effects affect the number of voxels associations with age. In Section 3.4, we present our experiments for simulating different levels of confounding between age and site. In Section 3.5 and Section 3.6, we present the consistency and replicability of the voxels associated with age for the different harmonization techniques. In Section 3.7, we present the bias in the associations between DTI values and age, and show how the different harmonization techniques perform at correcting for the bias. In Section 3.9, we show how the different harmonization techniques are robust to small sample size studies.

### 3.1 DTI scalar maps are highly affected by site

In Figure 4a), we present the histogram of FA values for the WM voxels for each participant, stratified by site. We observe a striking systematic difference between the two sites for all values of FA, with an overall difference of 0.07 in the WM (Welch two-sample t-test, *p* < 2.2e-16). Importantly, we notice that the inter-site variability in the histograms is much larger than the intra-site variability, confirming the importance of harmonizing the data across sites. A convenient way to visualize voxel-wise between-site differences in the FA values is plot the average between-site differences as a function of the average across sites. The Bland-Altman plot [Bland and Altman, 1986], also know as the Tukey mean-difference plot [Cleveland, 1993] or MA-plot [Dudoit et al., 2002, Bolstad et al., 2003] has been used extensively in the genomic literature to compare treatments and investigate dye bias. We use the more common terminology, MA-plot, and present the results for the FA values in Figure 4b). Not surprisingly, the scatterplot is shifted away from the zero line, indicating global site differences. Additionally, there is a large proportion of the voxels (top left voxels) appear to behave differently from other voxels. In the white matter atlas, these voxels are identified as being located in the occipital lobe (middle, inferior and superior gyri, and cuneus), in the fusiform gyrus and in the inferior temporal gyrus. This indicates that the site differences are region-specific, and that a global scaling approach will be most likely insufficient to correct for such local effects.

**Figure 4.**
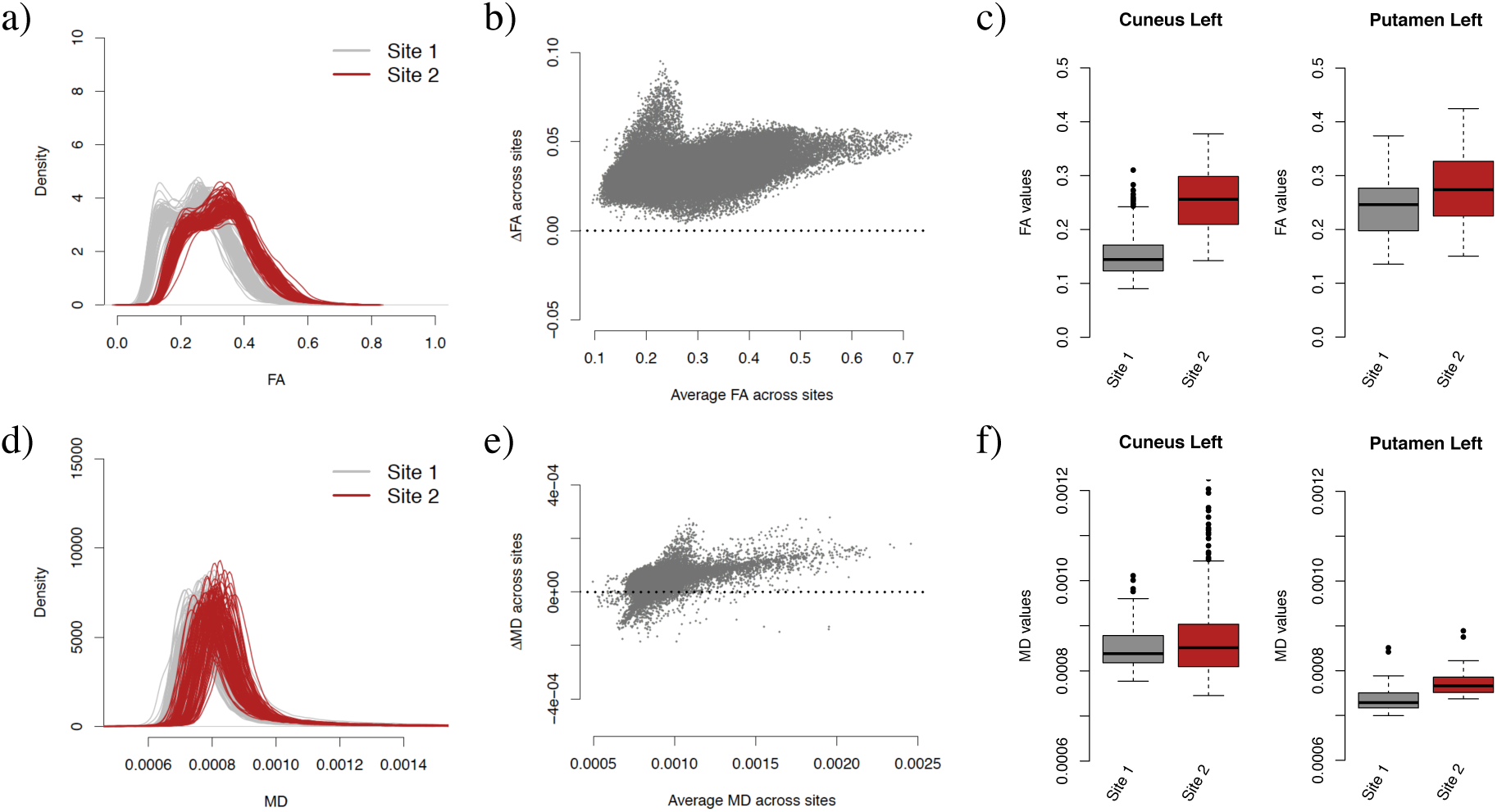
FA and MD maps are affected by site. **(a)** Density of the FA values for WM voxels for each participant, colored by site. **(b)** MA-plot for site differences in FA. The y-axis represents the differences in FA between Site 1 and Site 2, while the x-axis shows the average FA across sites. FA maps that would be free of site effects would result in an MA-plot centered around 0. The upper-left part of the scatterplot shows that several voxels appear to be differently affected by site in comparison to the rest of the voxels. **(c)** Boxplot of FA values for voxels located in two regions of interest (Cuneus left and Putamen left), depicted per site (FA values were averaged by site at each voxel separately). This shows that the magnitude of the difference in means between the two sites is region-specific. **(d-f)** Same as (a-c), but for the MD maps.

To further illustrate region-specific differences, we present in Figure 4c) the boxplots of FA values for two selected regions, cuneus left and putamen left, stratified by site. **Those results motivate the need of a region-specific harmonization**. We present similar results for the MD maps in Figure 4d,e,f. We note that the site differences appear to be more subtle for MD maps, but nonetheless present. Comparing panel c) and panel f), we observe that a brain region exhibiting site differences in FA maps do not necessarily show site differences in the MD maps.

### 3.2 ComBat successfully removes site effects in both FA and MD maps

In Figure 5, we present the MA-plots before and after each harmonization for the FA maps (see Figure A.3 for the MD maps). While both the scaling and Funnorm methods centered the MA-plots around 0, substantial technical variation remains, and local site-effects are still apparent. For the FA maps, RAVEL, SVA and ComBat reduce greatly the inter-site differences. For the MD maps, only SVA and ComBat successfully remove inter-site differences; RAVEL performs significantly worse. This is not surprising: in Figure A.1, it appears there is a lack of correlation between the average CSF value and average WM value in the MD maps. In other words, the CSF intensities do not act as site surrogates for the WM intensities, and therefore the RAVEL methodology underperforms in this situation.

Next, we calculated a t-statistic at each voxel to measure the association of the FA and MD values with site. We present in Figure 6a) the number of voxels in the WM that are significantly associated with site for each harmonization approach, for both FA and MD maps. A voxel is significant if the p-value calculated from the two-sample t-test is less than 0.05, after correcting for multiple comparisons using Bonferroni correction. Most voxels are associated with site in the absence of harmonization (raw data), and all harmonization methods reduce the number of voxels associated with site for both FA and MD maps at different degree. In agreement with the MA-plots, RAVEL, SVA and ComBat successfully remove site effects for most voxels in the FA maps, but only SVA and ComBat remove site effects for most voxels in the MD maps.

We also calculated t-statistics after summarizing FA and MD values by brain region. Using the Eve template atlas, we identified 156 region of interest (ROIs) overlapping with the WM mask. We present the number of regions significantly associated with site in Figure A.4a). While all ROIs are associated with site in the absence of harmonization in the FA maps, SVA and ComBat fully remove site effects for all ROIs. Residual site effects are found for more than a third of the ROIs for the Scaling, Funnorm and RAVEL harmonization methods. Similar results hold for the MD maps (140 out of 156 ROIs are affected by site in the absence of normalization).

**Figure 5.**
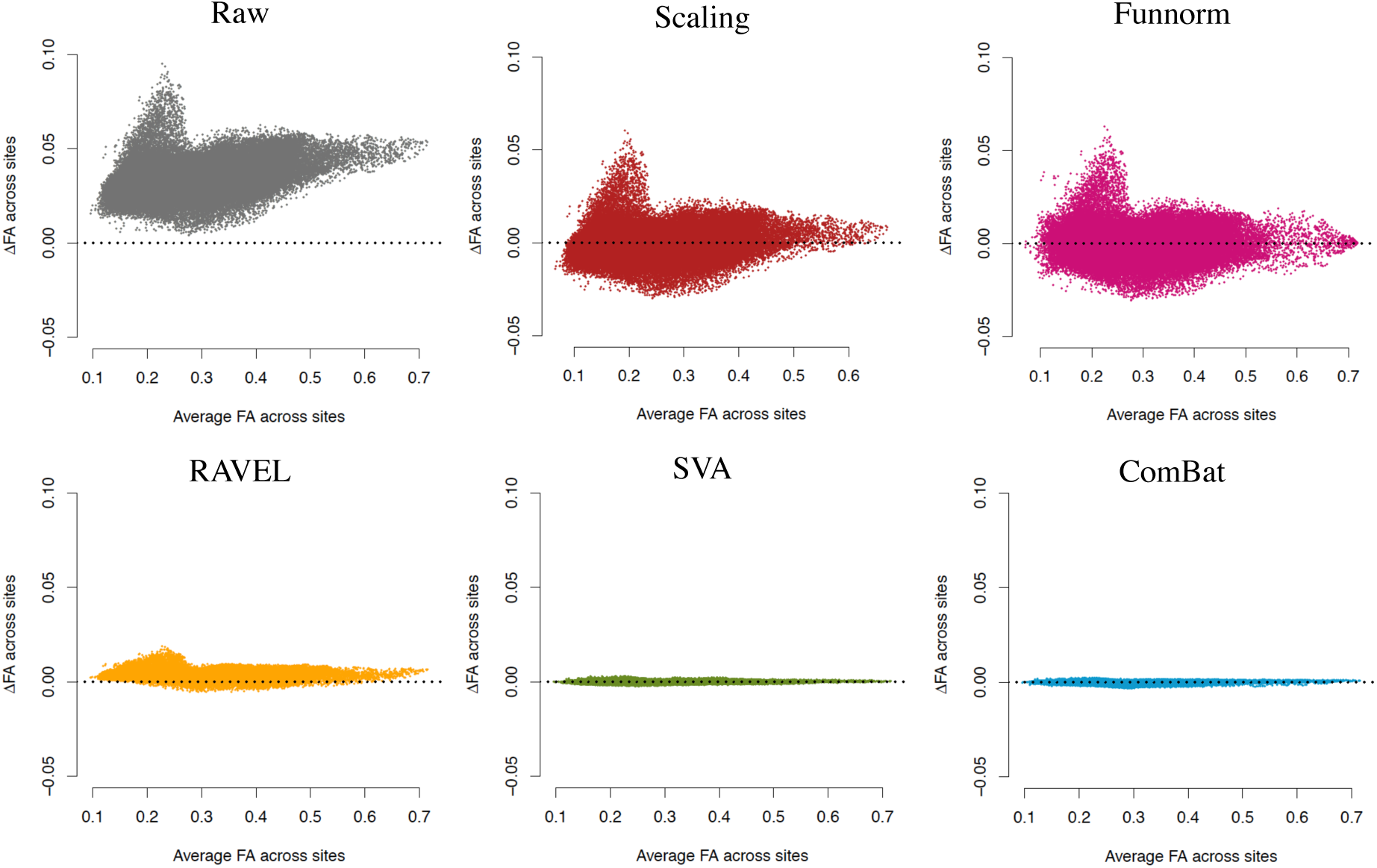
MA-plots for site differences in FA maps. Mean-difference (MA) plot for the FA maps for the different harmonization methods. At each voxel in the WM, the y-axis represents the difference between the average FA value at site 1 and the average FA value at site 2, and the x-axis represents the average FA value across all participants from both sites. An MA plot centered around 0 would indicate no global site differences. In the raw data, several voxels appear to be differently affected by site (upper-left points). The scaling and Funnorm methods successfully center the scatterplots around 0, but do not correct for local site effects. RAVEL, SVA and ComBat substantially perform better.

**Figure 6.**
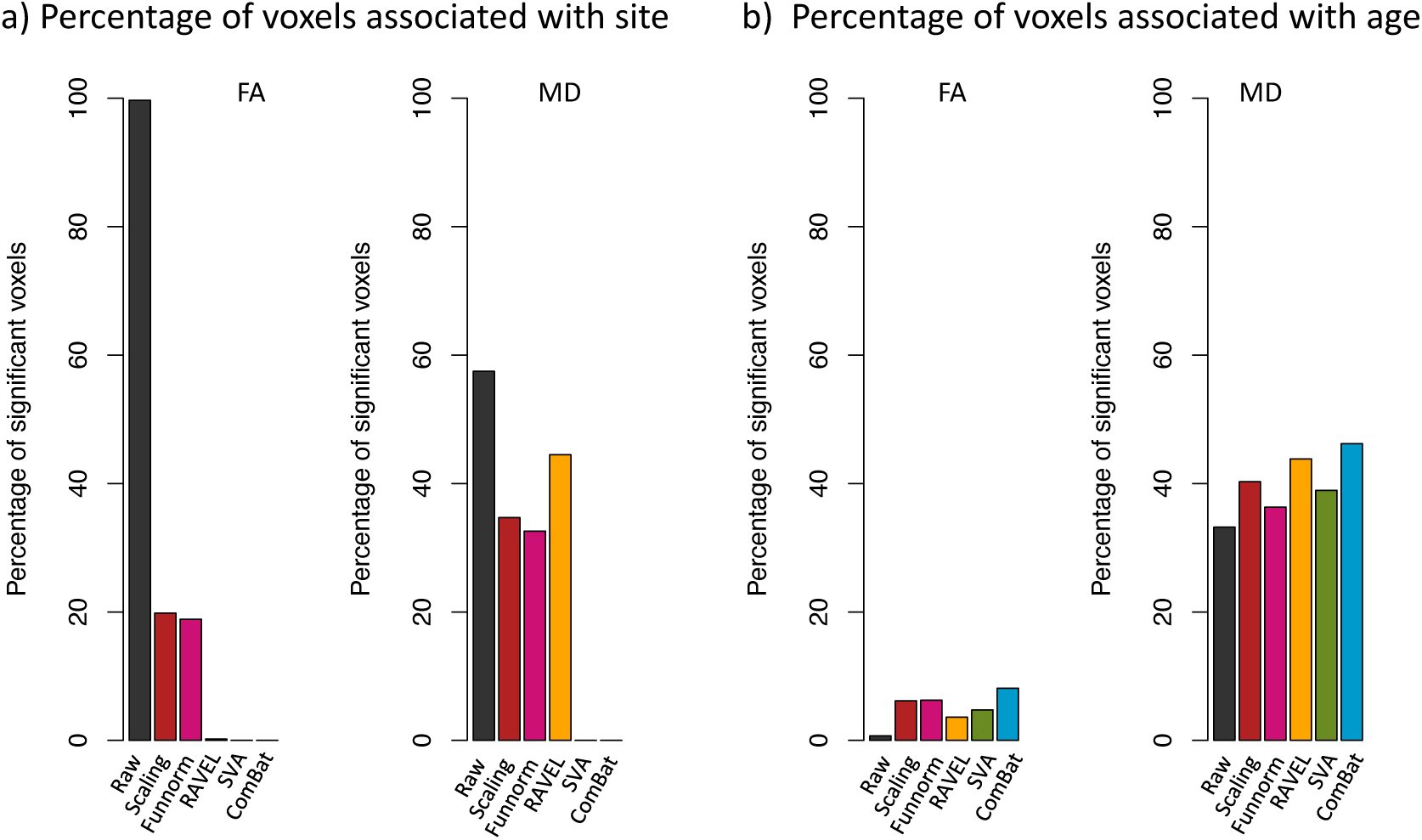
Number of voxels associated with site and age. **(a)** For each harmonization method, we calculated the number of voxels in the white matter (WM) that are significantly associated with imaging site for both FA and MD. A voxel is significant if the p-value calculated from a two-sample t-test is less than *p* < 0.05, after adjusting for multiple comparisons using Bonferroni correction. Lower numbers are desirable. **(b)** Number of voxels in the WM that are significantly associated with age using simple linear regression (*p* < 0.05) for both FA and MD. Higher numbers are desirable. From a total of 69,693 voxels in the WM, 69,475 and 40,056 voxels are associated with site in the raw data, for the FA and MD maps respectively. Both SVA and ComBat successfully remove the association with site for all voxels. ComBat performs the best at increasing the number of voxels associated with age (5,658 voxels for FA and 32,203 voxels for MD).

### 3.3 Harmonization across sites preserves within-site biological variability

A good harmonization technique should preserve the biological variability at each site separately. To test that, we calculated t-statistics for association with age before harmonization, for site 1 and site 2 separately, and after harmonization, for site 1 and site 2 separately as well. For each harmonization procedure, we computed the Spearman correlation between the unharmonized t-statistics and the harmonized statistics. For Site 1, the correlations are: ρ =0.997 for both Scaling and Funnorm, *ρ* = 0.981 for RAVEL, *ρ* = 0.893 for SVA and *ρ* = 0.994 for ComBat. For Site 2, the correlations are: *ρ* = 0.996 for both Scaling and Funnorm, *ρ* = 0.964 for RAVEL, *ρ* = 0.875 for SVA and *ρ* = 0.997 for ComBat. The ComBat, Scaling and Funnorm methods perform exceptionally well. We note that the correlation is substantially lower for SVA at both sites. This is not surprising; unlike other methods, SVA removes variability that is not associated with age across the whole dataset, but does not protect for the removal of biological variability at each site individually.

In Figure (6b), we present the number of voxels in the WM that are significantly associated with age for each harmonization approach, for both FA and MD maps. A voxel is called significant if the p-value calculated from simple linear regression is less than 0.05, after adjusting for multiple comparisons using Bonferroni correction. All harmonization methods increase the number of significant voxels associated with age in comparison to the raw data. ComBat presents the most substantial gain for FA maps (5658 voxels, in comparison to 481 voxels for raw data) and for MD maps (32,203 voxels, in comparison to 23,136 voxels for raw data). We also performed a similar analysis at the ROI level: using the white matter atlas, we computed an average FA value at each region, for each participant separately, and subsequently applied the different harmonization techniques; similar results were obtained (see Figure A.4b in Appendix).

### 3.4 Harmonization and confounding

In the next sections, we evaluate the performance of the different harmonization procedures by estimating the consistency and replicability of the voxels associated with age. We also investigate the robustness of the different harmonization techniques to datasets for which age is confounded by site. The previous results were obtained by harmonizing two sites that were carefully matched for age, gender and ethnicity to minimize potential confounding of those variables with site. However, matching has several limitations. If there is a poor overlap between the covariates of interest across sites, matching will result in a significant exclusion of samples. In addition, the number of scans to be excluded is proportional to the number of covariates to be matched, which can be significant in many applications, making matching infeasible. On the other hand, failing to match for covariates will result in an undesirable situation where site will be a confounder for the relationship between the DTI values and the phenotype(s) of interest. Thus, a better alternative to matching is to first combine all available data across sites, and then to apply a harmonization technique that is robust to confounding.

Confounding between age and site presents an additional challenge for harmonization, since removing variation associated site can lead to removing variation associated with age if not done carefully. To evaluate the robustness of the different harmonization methods in the presence of statistical confounding between imaging site and age (that is age is unbalanced with respect to site), we selected different subsets of the data to create several confounding scenarios, as shown in Figure 7. For illustration purpose, we chose a voxel in the right thalamus for which the association between FA and age is high. Figure A.2 illustrates the confounding scenarios using median FA values in the WM. We see that for the full data (Figure 7a), the FA values increase linearly with age within each site.

**Figure 7.**
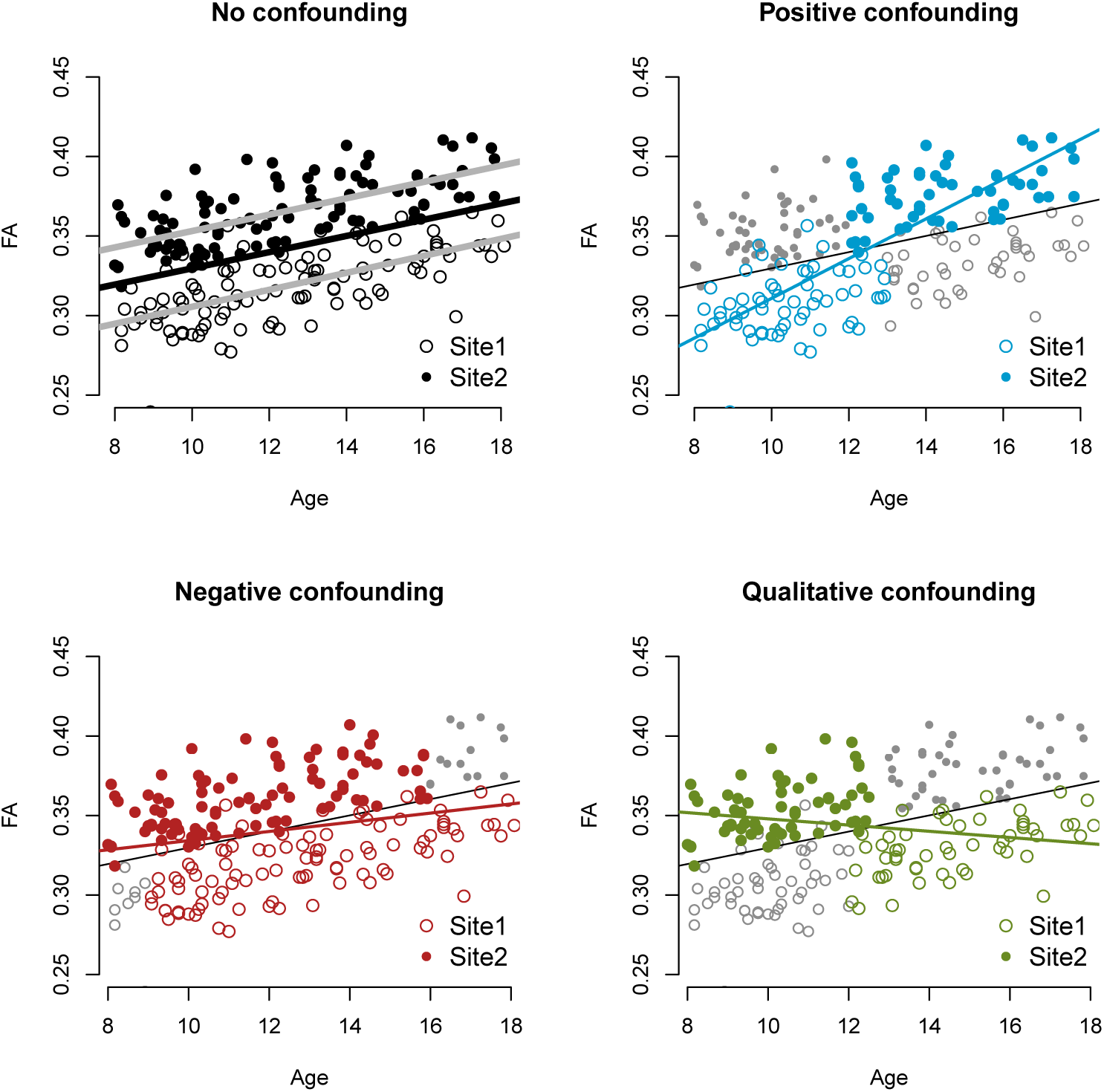
Confounding scenarios for FA maps. In all four panels, each data point represents the FA value versus the age of the participant for a fixed voxel in the right thalamus. Full dots and circles are used to distinguish the two sites of the participant scans (Dataset 1 and Dataset 2). The solid black line in all panels represents the estimated linear relationship between FA and age when all data points are included (absence of confounding). In panel (a), the grey lines represent the estimated relationship between FA and age for each site. In panels (b-d), the selected participants are colored (blue, red and green respectively), and the colored solid lines represent the estimated linear relationship between FA and age for the selected participants only.

“Positive confounding” and “negative confounding” refer to situations where the relationship between the FA values and age is overestimated and underestimated, respectively, with the same directionality of the true effect. Selecting older samples from Site 2, and younger samples from Site 1, creates positive confounding (Figure 7b). This is because the average FA value for Site 2 is higher than the average value for Site 1. On the other hand, excluding the oldest participants from Site 2 and the youngest participants from Site 1 will create negative confounding (Figure 7c). “Qualitative confounding” is an extreme case of confounding where the estimated direction of the association is reversed with respect to the true association. Selecting younger participants from Site 2, and older participants from Site 1, with no overlap of age between the two sites, creates such confounding (Figure 7d).

We note that in the no-confounding scenario of Figure 7a), the association between the FA values is unbiased in the sense that it is not modified by site. Indeed, the slope using all the data (black line) is similar to the slopes estimated within each site (grey lines). However, the variance of the estimated slope will be inflated due to the unaccounted variation attributable to site.

We now present the consistency and replicability of the voxels associated with age for the different harmonization methods in the different confounding scenarios.

### 3.5 ComBat improves the consistency of the voxels associated with age

We evaluated the consistency of the voxels associated with age using the discovery-validation scheme described in Section 2.4.1. We considered the harmonized dataset as a discovery cohort, and the two within-site datasets as validation cohorts. We performed a mass-univariate analysis testing for association with age separately for each cohort, and used CAT curves [Irizarry et al., 2005] to measure the consistency of the results between the discovery and validation cohorts.

We present in Figure 8 the CAT curves for each of the confounding situation. In the absence of confounding (first column), there is good overlap for all methods, including the raw data. ComBat performs the best, with a flat CAT curve around 1. This indicates that the ranking of the t-statistics in the ComBat-harmonized dataset is almost the same as the ranking of the t-statistics estimated at each site separately. In the positive confounding scenario (second column), all methods perform similar to the raw data, except for the scaling and Funnorm approaches that show substantial inconsistencies with the ranking of the within-site t-statistics, as seen by their CAT curves closer to the diagonal line. This is not surprising; both the scaling and Funnorm approach are global approaches, and because of the positive nature of the confounding, the removal of a global shift associated with site will also remove the global signal associated with age.

**Figure 8.**
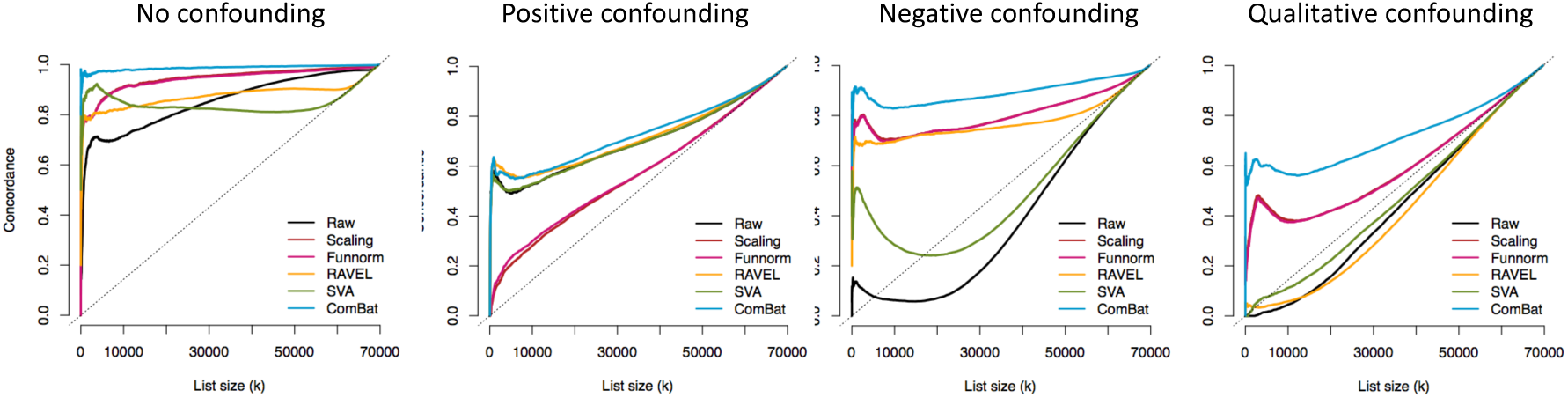
Consistency of the voxels associated with age in the FA maps. For each confounding scenario and for each harmonization method, we calculated a concordance at the top (CAT) curve for voxels associated with age. The concordances were calculated between the harmonized dataset (2 sites combined) and each of the two within-site datasets; the CAT curves presented here are averaged across the two within-site datasets. Briefly, for each value of the x-axis (*k*), the CAT curve measures the proportion of the top *k* voxels associated with age in the harmonized dataset that are also present in the top *k* voxels associated with age for each of the within-site dataset. The voxels associated with age from the two within-site datasets represent gold-standard voxels. A good harmonization will increase the overlap with those voxels and will result in a CAT curve closer to 1. Overlaps by chance will result in a CAT curve along the diagonal dashed line indicating change agreement.

In the presence of negative confounding and qualitative confounding, combining the data without a proper harmonization technique lead to more severe problems (Figure 8, third and fourth columns). The CAT curves for the raw data (no harmonization) are considerably below the diagonal line, indicating a negative correlation between the within-site t-statistics ranking and the combined dataset t-statistics ranking. The negative correlation can be explained by the following: because of the negative (or qualitative) confounding, the t-statistics for the voxels that are truly not associated with age, normally centered around 0, became highly negative because of the site effect. On the other hand, the t-statistics for the voxels associated with age, normally positive for FA, are shifted towards 0. The negative and qualitative confounding artificially make the null voxels significant and create a reversed ranking. In the negative confounding scenario, all methods, except SVA, are able to recover a ranking that is much more consistent with the true ranking, therefore improving consistency of the results. In the qualitative confounding situation, only ComBat, Funnorm and the scaling approach improving upon the raw data, with ComBat showing the greatest improvement. Overall, the results are promising for ComBat: the consistency of the top voxels associated with age is dramatically improved for all confounding scenarios, making ComBat a robust harmonization method. The other harmonization approaches show variable performance.

### 3.6 ComBat improves the replicability of the voxels associated with age

The CAT curves presented in the previous section were calculated using the two site datasets as validation cohorts, which are not independent with respect to the harmonized dataset. In this section, we calculate the CAT curves using an independent dataset as validation cohort. This evaluates the performance of the different harmonization techniques at replicating the voxels associated with age across independent datasets, where replicability is defined as chance that an independent experiment will produce consistent results [Leek and Peng, 2015]. We note that replicability is a stronger result than consistency since it evaluates the methods using external validation.

We used two different independent cohorts for estimating the replicability: a larger cohort composed of 292 participants with a similar age distribution (Independent Dataset 1) and a cohort composed of 105 participants with a slightly older age distribution (Independent Dataset 2), as described in the Methods section. Both cohorts were taken from the PNC [Satterthwaite et al., 2014].

In Figure 9a), we present the CAT curves using Independent Dataset 1 as a validation cohort (same age range). The results are very similar to the consistency results presented in Figure 8. ComBat performs the best at replicating the voxels associated with age for all confounding scenarios. In Figure 9b), we present the CAT curves using Independent Dataset 2 as a validation cohort (older age range). Because the population of the independent cohort in this case is older, there may be differences in the subset of voxels that are truly associated with age. This can be seen in lower overall concordances curves in Figure 9b). Nevertheless, ComBat still performs the best at improving the replicability of the voxels, for all confounding scenarios.

**Figure 9.**
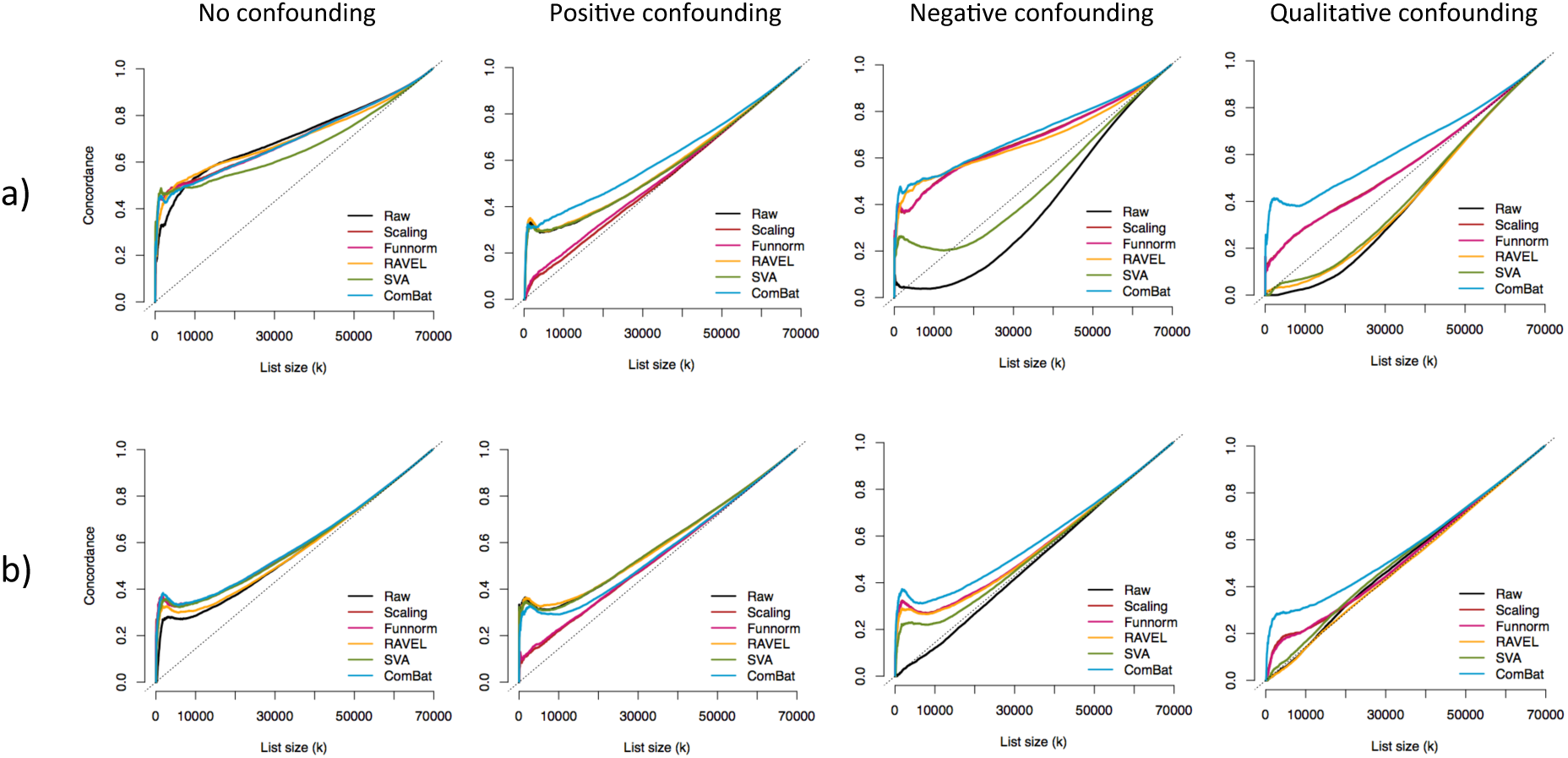
Replicability of the voxels associated with age in the FA maps. For each confounding scenario and for each harmonization method, we calculated a concordance at the top (CAT) curve for the voxels associated with age. The concordances were calculated between the harmonized dataset (2 sites combined) and an independent dataset. In **(a)**, 292 unrelated participants within the same range were selected as an independent cohort. In **(b)**, 105 unrelated and older participants were selected as an independent cohort. A good harmonization will result in a CAT curve closer to 1. Overlaps by chance will result in a CAT curve along the diagonal.

### 3.7 ComBat successfully recovers the true effect sizes

In this section, we evaluate the bias in the estimated changes in FA associated with age 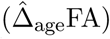 for each harmonization procedure, for the different confounding scenarios. We refer to 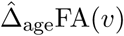) as the estimated “effect size” for voxel *v*. The effect site can be estimated using linear regression (slope coefficient associated with age). In principle, to assess unbiasedness, we would need to know the true effect sizes Δ_age_FA. We circumvent this by estimating the effect sizes on the signal silver-standard described in Section 2.4.3. For each site, we calculated the effect size for each voxel of the signal silver-standard by running a simple linear regression for age, and retaining the regression coefficient for age as the estimated effect size. We took the average across the two sites at each voxel as the estimated true effect size. This resulted in a distribution of 2265 effect sizes for the signal voxels, with a median effect size close to 0.004, presented in the left boxplot of Figure A.6a). We also estimated the true effect sizes for voxels not associated with age (null voxels described in Section 2.4.3). We obtained a distribution of 1932 effect sizes for the null signal. Not surprisingly, those effects sizes are roughly centered at 0 (right boxplot, Figure A.6a).

In Figure 10a), we present the distribution of the estimated effect sizes on the signal silver-standard for all methods, and for all confounding scenarios. The dashed lined represents the median effect size of the true effect sizes, and the solid line represents an effect size of 0. As expected, the effect sizes in the raw data (datasets combined without harmonization) are consistent with the type of confounding; positive confounding shifts the effect sizes positively, and the negative and qualitative confounding shifts the effect sizes negatively. ComBat is the only harmonization technique that fully recovers the true effect sizes for all confounding scenarios in terms of median value and variability. Funnorm and RAVEL both reduced the bias in the effect sizes distribution, and both underestimate the true associations. We note that RAVEL performs sensibly worse for the qualitative confounding scenario. Interestingly, SVA does not achieve any bias correction for any of the confounding scenarios; the distribution of the estimated effect sizes resemble those of the unharmonized dataset. This could be explained by the fact that SVA method “protects” for the present association between the outcome and the covariate of interest, and therefore an association that is biased in the original dataset will remain biased in the SVA-corrected dataset. Similarly, we present in Figure 10b) the distribution of the estimated effect sizes for the null silver-standard. We recall that a successful harmonization approach will result in a boxplot centered around 0. The results are similar to Figure 10a); ComBat successfully recovers the true effect size distribution for all confounding scenarios. Results for MD maps are presented in Figure A.6b) and Figure A.7.

**Figure 10.**
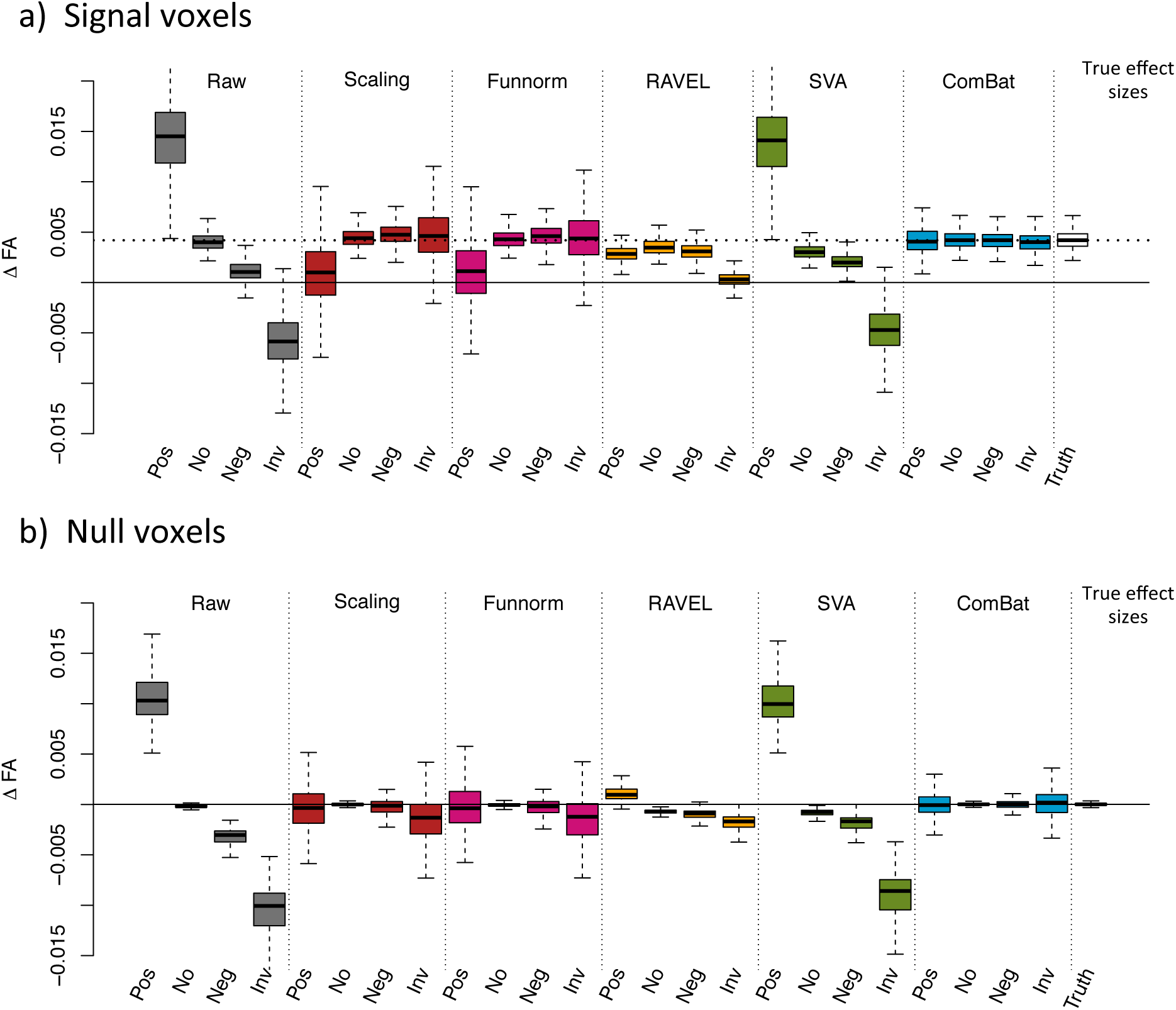
Estimated effect 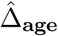FA for different confounding scenarios. **(a)** Boxplots of the estimated effect sizes 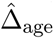FA for the set of signal voxels described in Section 3.9 for different confounding scenarios: positive confounding (pos), no confounding (no), negative confounding (neg) and quantitative confounding (rev). The dotted line represents the median true effect size (around 0.004). **(b)** Boxplots of the estimated effect sizes 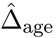 for the set of null voxels described in Section 3.9. The median true effect size is around 0. The distributions of the estimated effect sizes for the ComBat-harmonized datasets approximate very well the distribution of the true effect sizes shown in the last column in each panel. Results for MD values are presented in Figure A.7.

The retrieval of unbiased effect sizes for both the signal and the null silver-standard strongly suggests that ComBat successfully removed the site effect in the combined datasets without removing the signal associated with age, even in the presence of substantial confounding between age and site. The FA changes estimated after ComBat for voxels highly associated with age are similar to the FA changes measured at each site separately.

### 3.8 ComBat improves statistical power

In Figure 11, we present the distribution of the WM voxels-wise t-statistics measuring association with age in the FA maps for four combinations of the data: Site 1 and Site 2 analyzed separately, Site 1 and Site 2 combined without harmonization, and Site 1 and Site 2 combined and harmonized with ComBat. The goal of combining datasets from different sites is to increase the sample size, and therefore the power of the statistical analysis. We therefore expect t-statistics with higher magnitude for voxels truly associated with age. Moreover, we note that most of the t-statistics will be positive as a consequence of the global increase in FA associated with development of the brain in teenagers [Tamnes et al., 2010, Bava et al., 2010, Lebel and Beaulieu, 2011].

**Figure 11.**
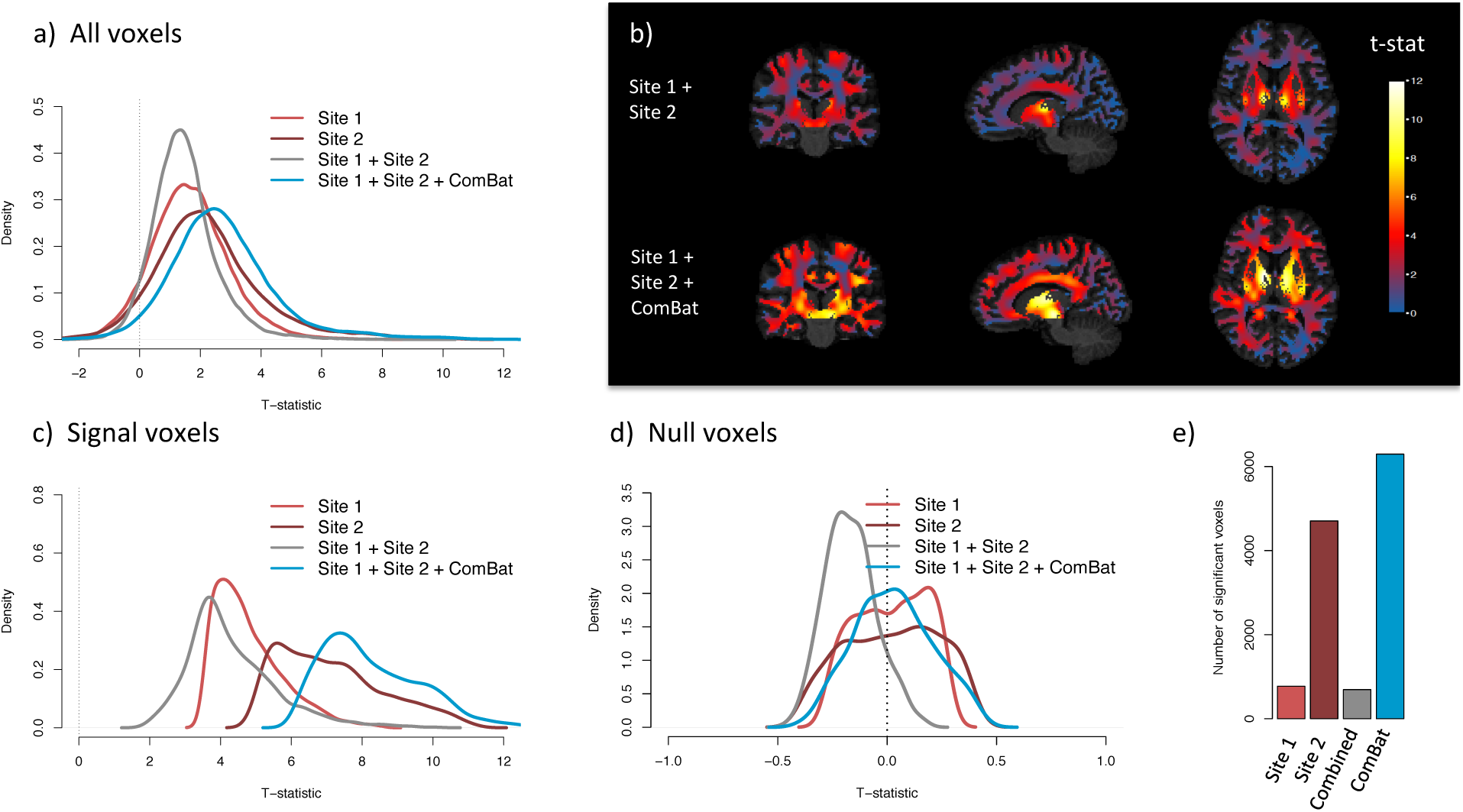
ComBat improves statistical power. We present voxel-wise t-statistics in the WM, testing for association between FA values and age, for four combinations of the data: Dataset 1 and Dataset 2 analyzed separately, Dataset 1 and Dataset 2 combined without any harmonization, and Dataset 1 and Dataset 2 combined and harmonized with ComBat. **(a)** Distribution of the t-statistics for all WM voxels, for each analyzed dataset. The combined datasets harmonized with ComBat show higher t-statistics. **(b)** T-statistics in template space for the combined dataset, with no harmonization (top row) and with Combat (bottom row). **(c)** Distribution of the t-statistics for a subset of voxels highly associated with age (signal silver-standard described in Section 2.4.3). **(d)** Distribution of the t-statistics for a set of voxels not associated with age (null silver-standard described in Section 2.4.3). ComBat increases the magnitude of the t-statistics for the signal voxels while maintaining the t-statistics around 0 for the null voxels. **(e)** Number of voxels significantly associated with age. Bonferroni correction was applied to correct for multiple comparisons

In Figure 11a), in which we present the t-statistics for all voxels in the WM, we observe an opposite effect. The distribution of the t-statistics for the two sites combined without harmonization is shifted towards 0 (mean t-statistic of 1.4) in comparison to the t-statistics obtained from both sites separately (mean t-statistic of 1.7 and 2.3 for site 1 and site 2 respectively).This strongly indicates that combining data from multiple sites, without harmonization, is counter-productive and impairs the quality of the data. On the other hand, combining and harmonizing data with ComBat results in a distribution of higher t-statistics on average (mean t-statistic of 2.8). We present in Figure 11b) the t-statistics in template space with and without ComBat.

To further examine the effects of harmonization on the data, we present the distribution of the t-statistics for voxels that are truly associated with age (signal silver-standard described in Section 2.4.3) in Figure 11c, and voxels that are truly not associated with age (null silver-standard described in Section 2.4.3) in Figure 11d). This confirms that ComBat increases the statistical power at finding voxels truly associated with age, as seen by the distribution of t-statistics substantially shifted to the right in Figure 11c. The mean t-statistic for the raw data and after ComBat is 4.3 and 8.3 respectively. ComBat also keeps the t-statistics of the null voxels tightly centered around 0 (Figure 11d). In Figure 11e), we present the number of voxels significantly associated with age (*p* < 0.05) after adjusting for multiple comparisons using Bonferroni correction. The results strengthen our observations that harmonization is needed in order to successfully combine multi-site data.

We present the results for the MD maps in Figure A.5. It is expected to observe many voxels showing a negative association between MD and age in teenagers [Tamnes et al., 2010, Bava et al., 2010, Lebel and Beaulieu, 2011], and therefore to observe a distribution of t-statistics shifted towards negative values (as opposed to the t-statistics distribution in FA maps). Again, ComBat successfully increases the magnitude of the t-statistics for the signal voxels (distribution of the t-statistics highly shifted away from 0 in Figure A.5c), while maintaining the t-statistics for the null voxels centered around 0 (Figure A.5d).

### 3.9 ComBat is robust to small sample sizes

A major advantage of ComBat over other methods is the use of Empirical Bayes to improve the estimation and removal of the site effects in small sample size settings. To assess the robustness of the different harmonization approaches for combining small samples size studies, we created *B* = 100 random subsets of size *n* = 20 across sites. Specifically, we selected for each subset 10 participants at random from each site. For each subset, we applied the different harmonization methods and calculated voxel-wise t-statistics in the WM, for testing the association of the FA values with age, for a total of 100 t-statistic maps. To obtain an estimated gold-standard for a t-statistic map obtained with studies of sample size 20, that we refer to as a silver-standard, we created *B* = 100 random subsets of size 20 from site 1, and *B* = 100 additional random subsets of size 20 from site 2. Because subsets are created within site, they are not affected by site effects and results obtained from those subsets should be superior or as good as any of the results obtained from the harmonized subsets.

In Figure 12a), we present the average CAT curve for each harmonization method (average taken across the random subsets) together with the silver-standard CAT curve (dark blue), for the FA maps. All methods improve the replicability of the voxels associated with age. We note that Combat performs as well as the silver-standard, successfully removing most of the site effects. In Figure 12b), we present the densities of the t-statistics for the top voxels associated with age (signal voxels described in Section 2.4.3) for the FA maps. We note that all methods improve the magnitude of the t-statistics, therefore increasing statistical power, with ComBat showing the best performance, notably performing as well as the silver-standard. In Figure 12c), we present the densities of the t-statistics for voxels not associated with age (null voxels described in Section 2.4.3) for the FA maps; a good harmonization method should result in t-statistics centered around 0. The global scaling approach, functional normalization and ComBat correctly correctly return t-statistics centered around 0 that are similar to the silver-standard. SVA and RAVEL do not perform as well (densities shifted away from 0). Overall, the results show that ComBat is a very promising harmonization method even for small sample size studies, doing as well as a dataset that was not affected by site effects. Similar results were obtained for the MD maps, presented in the panels (d-f) of Figure 12.

**Figure 12.**
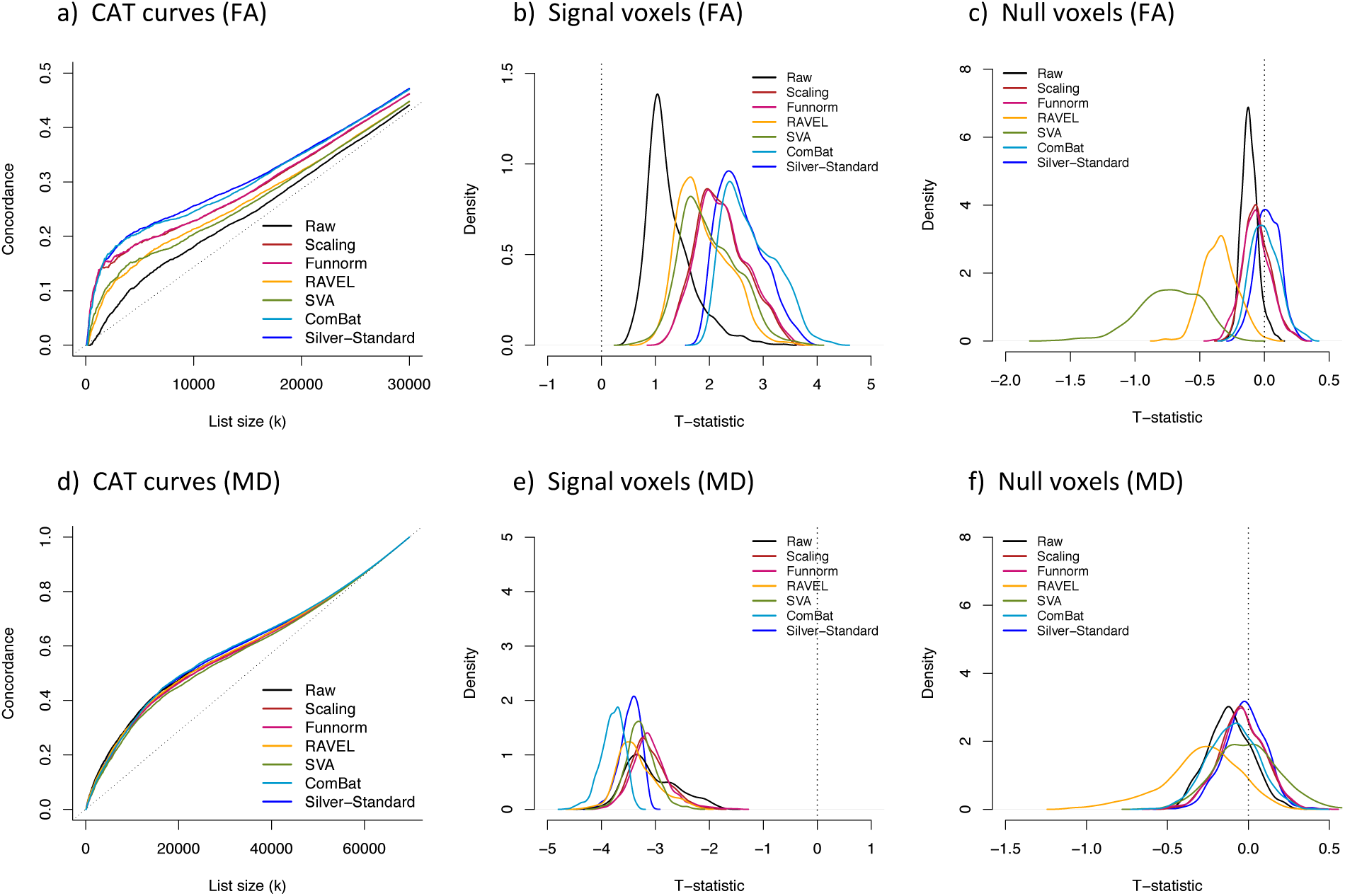
ComBat is robust to small sample size studies. We created *B* = 100 random subsets of size 20, selecting at random 10 participants from each site, and applied each harmonization method on every subset separately. For each harmonized subset, we computed a t-statistic at each voxel in the WM, testing for the association of FA and MD with age. We created a silver-standard list of t-statistics by creating *B* = 100 random subsets of size 20 within site. **(a)** Average concordance at the top (CAT) curve for each harmonization method for the FA maps. The silver-standard CAT curve is depicted in dark blue. A higher curve represents better replicability of the voxels associated with age. **(b)** Densities of the t-statistics for the set of signal voxels described in Section 3.9, for the FA maps. Higher values of the t-statistics are desirable. **(c)** Densities of the t-statistics for the set of null voxels described in Section 3.9, for the FA maps. T-statistics closer to 0 are desirable. For each plot, the results obtained for the ComBat-harmonized datasets approximate very well the results obtained from the within-site silver-standard (dark blue). **(d)** Same as **(a)**, but for the MD maps. **(e)** Same as **(b)**, but for the MD maps. Lower values of the t-statistics are desirable. **(f)** Same as **(c)**, but for the MD maps. RAVEL performs substantially worse than other methods.

## 4 Discussion

In this work, we investigated the effects of combining DTI studies across sites and scanners on the statistical analyses. We used FA and MD maps from data acquired at two sites with different scanners. We first showed that combining the two studies without proper harmonization led to a decrease in power of detecting voxels associated with age. This confirmed that DTI measurements are highly affected by small changes in the scanner parameters, as those affect the underlying water diffusivity. This motivated the need for harmonizing data across sites and scanners. We then adapted and compared several statistical harmonization techniques for DTI studies.

Using a comprehensive evaluation framework that respects the importance of biological variation in the data, we showed that ComBat, a popular batch effect correction tool used in genomics, performs the best at harmonizing FA and MD maps. It allows site effects to be location-specific, but pools information across voxels to improve the statistical estimation of the site effects. More specifically, we showed that ComBat substantially increases the replicability of the voxels associated with age across independent experiments. We also investigated the robustness of the proposed harmonization methods when the associations of age and DTI measurements are confounded by site as a consequence of possible unbalanced data, as well as robustness to small sample sizes. ComBat was the best at improving the results across all scenarios, and appeared to be robust to small sample size studies. Indeed, it was able to recover the true associations between the FA (and MD) values and age, despite the bias introduced by the association between site and age.

Global scaling and functional normalization [Fortin et al., 2014], did not perform well overall. This is not surprising; both of these histogram-normalization methods fail to account for the spatial heterogeneity of the site effects throughout the brain. We also compared ComBat to RAVEL, an intensity normalization technique previously proposed for T1-w images [Fortin et al., 2016a]. RAVEL performed well for the FA maps, for which the FA values in the CSF reflect the technical variation in the WM. However, RAVEL did not perform well for the MD maps; the site effects in the CSF were not correlated with the site effects in the WM. We also compared ComBat to SVA [Leek and Storey, 2007, 2008], an algorithm developed for genomics data that estimates unwanted variation that is orthogonal to the biological variation. SVA was successful at estimating and removing the site effects, but did not perform as well as ComBat for datasets for which age was confounded with site.

The ComBat methodology can be extended in several ways. In the case of a categorical outcome, for instance disease status or gender, a first extension to ComBat would be to estimate the voxel-specific site effect parameters only using participants from a reference category. This would be particularly useful for datasets with unbalanced data, as demonstrated in Linn et al. [2016b]. Future ComBat models might also draw strength from spatial correlation by spatially restraining the estimation of the hyperparameters for the prior distributions to only pool information across neighboring voxels. Another extension would be to incorporate an inverse probability weighting (IPW) scheme to explicitly model statistical confounding between the phenotype of interest in site. IPW has been shown to improve results when there is presence of confounding in imaging studies [Linn et al., 2016a], especially in mitigating multivariate confounding for prediction.

While this paper has focused on the harmonization of imaging data across sites and scanners, another important challenge is the harmonization of imaging data within a site. Indeed, even for scans acquired on the same scanner, between-participant unwanted variation that is technical in nature also exists. This requires a harmonization technique that is not dependent on a site, or scanner variable. In genomics, latent factor approaches that estimate unknown source of variation have been successfully used, such as SVA [Leek and Storey, 2007] and RUV [Gagnon-Bartsch and Speed, 2012]. Recently, a similar approach called RAVEL [Fortin et al., 2016a] has been developed for the harmonization of structural MRI intensities using a control region for the estimation of the latent factors of unwanted variation [Fortin et al., 2016a]. The choice such a control region can be difficult depending on the phenotype and modalities of interest.

Although we have shown the performance of ComBat in the context of DTI scalar maps, the ComBat model is widely applicable beyond this setting. It can also be used to harmonize connectivity data across different processing protocols, such as seed-based connectivity maps in resting-state fMRI or measures of structural connectivity derived from DTI. In addition, while we used voxels as features to be harmonized in the ComBat model, the ComBat technique can be applied to measurements summarized at the ROI level, making ComBat a promising harmonization technique for other imaging modalities, including for instance volumetric and cortical thickness studies.

## 5 Software and reproducible analysis

All of the postprocessing analysis was performed in the R statistical software (version 3.2.0). For SVA and ComBat, reference implementations from the sva package were used. All figures were generated in R with customized and reproducible scripts, using several functions from the package fslr [Muschelli et al., 2015]. We have adapted and implemented the ComBat methodology to imaging data, and the software is available in an R package on GitHub (https://github.com/Jfortin1/ComBatHarmonization).

## Abbreviations

ADNI: Alzheimer’s Disease NeuroImaging Initiative
AX: Axial diffusivity
CAT: Concordance at the top
ComBat: Combatting batch effects when combining batches of gene expression microar-ray data
CoV: Coefficient of variation
CSF: Cerebrospinal fluid
DTI: Diffusion tensor imaging
EB: Empirical Bayes
FA: Fractional anisotropy
GM: Grey matter
GS: Global scaling
IBMA: Image-based meta analysis
IPW: Inverse probability weighting
MD: Mean diffusivity
MRI: Magnetic resonance imaging
OLS: Ordinary least squares
RAD: Radial diffusivity
RAVEL: Removal of artificial voxel effect by linear regression
RISH: Rotation invariant spherical harmonic
ROI: Region of interest
SVA: Surrogate variable analysis
SVD: Singular value decomposition T1-
w: T1-weighted
TBSS: Tract-based spatial statistics
WM: White matter
WMPM: White matter parcellation map;

## Authors contributions

JPF developed the methodology and analyzed the data. DP, BT and TW processed the data. ME, KR, DR, TS, RCG, REG and RTSc recruited the participants and acquired the data. JPF and RTSh wrote the manuscript. RTSh and RV supervised the work. All authors read and approved the final manuscript.

## Appendix

**Figure A.1.**
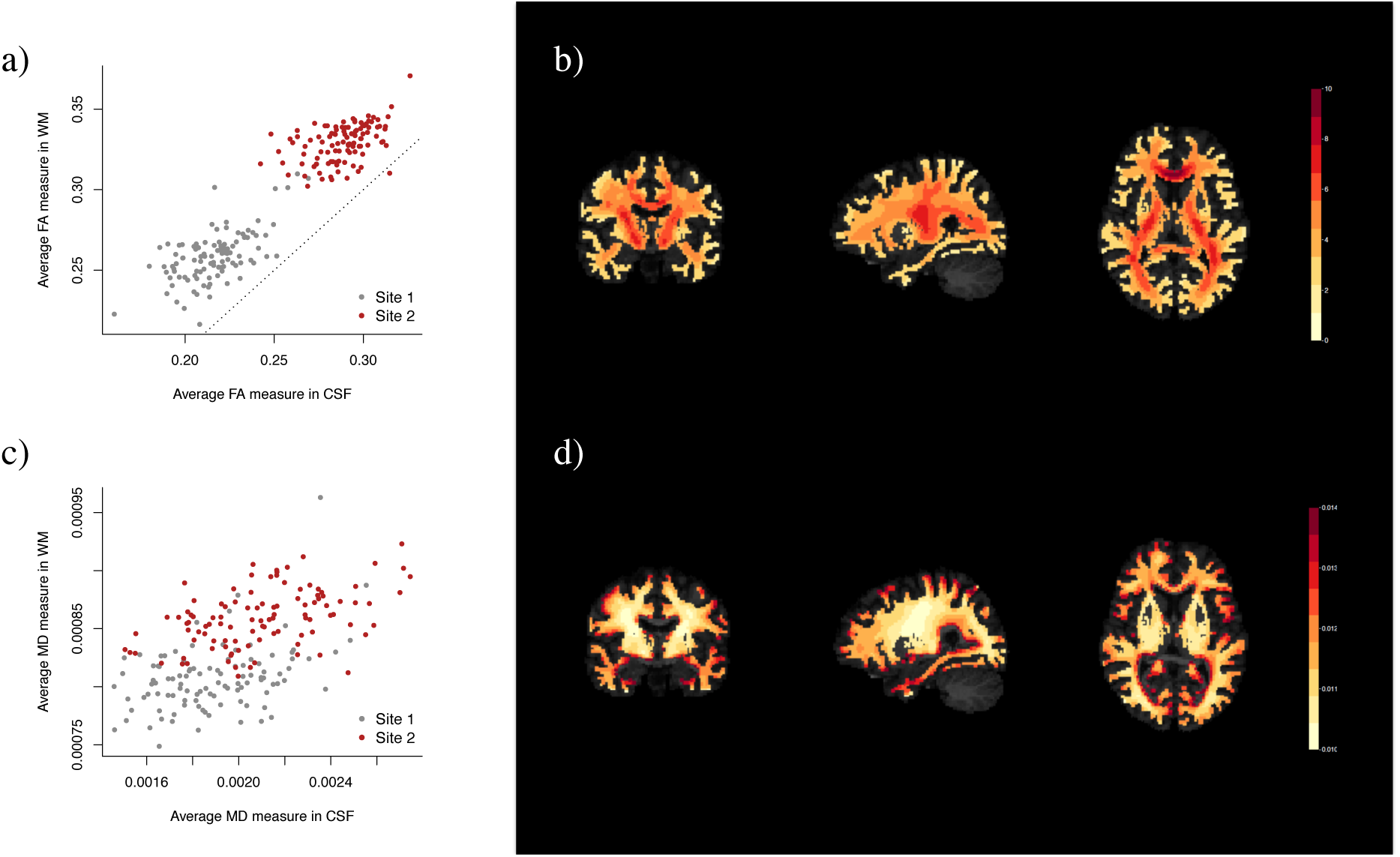
RAVEL Aharmonization. **(a)** Relationship between the average FA measure in white matter (WM) and cerebrospinal fluid (CSF). The FA measurements vary by site in both WM and CSF. **(b)** Voxel-specific RAVEL coefficient 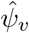 in template space for FA maps. **(c)** Relationship between the average MD measure in white matter (WM) and cerebrospinal fluid (CSF). The MD measurements vary by site in WM, but do not seem to vary in CSF. **(d)** Voxel-specific RAVEL coefficient 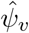 in template space for MD maps.

**Figure A.2.**
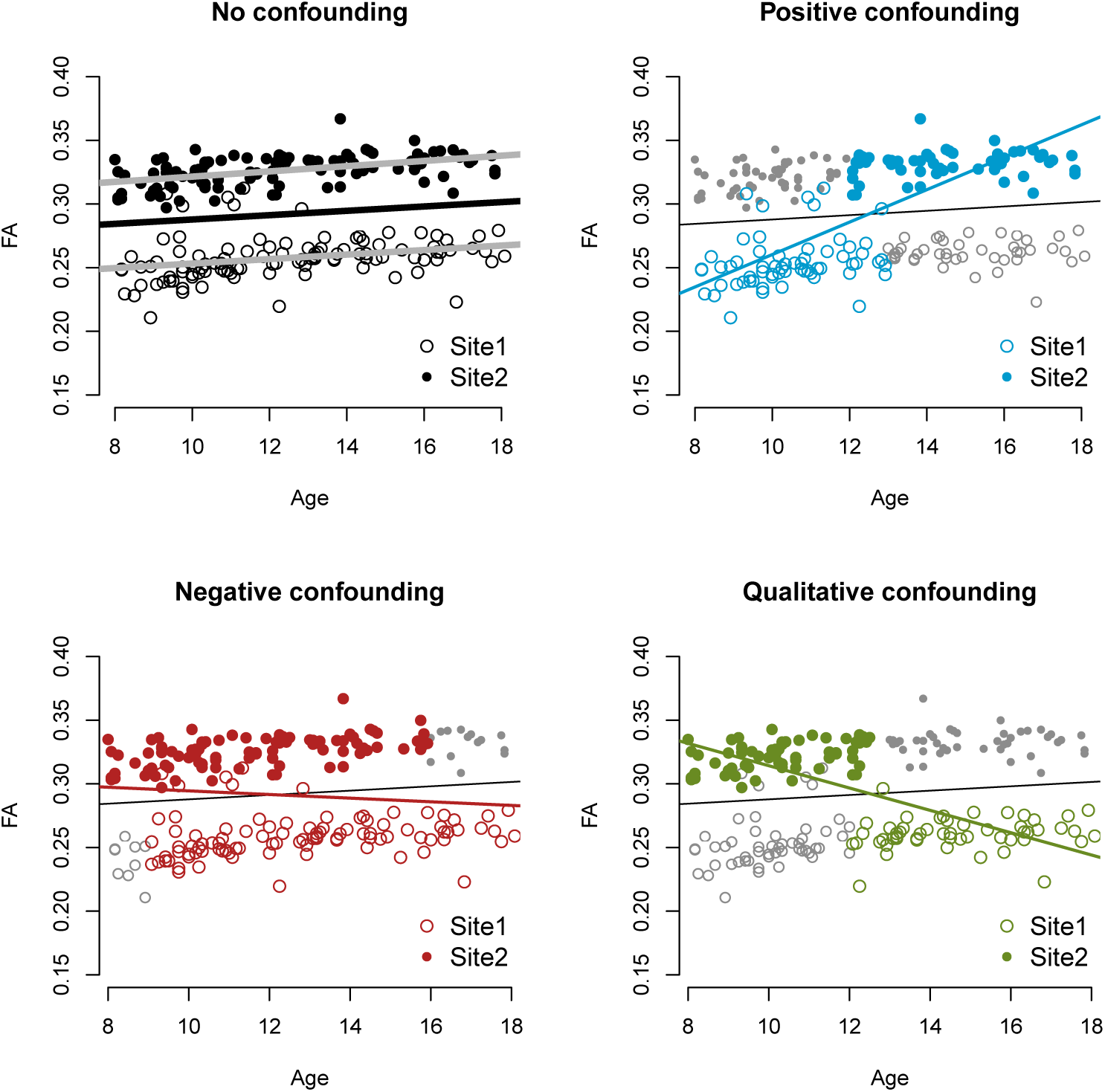
Confounding scenarios for FA maps. Same as Figure 7, but for the per-scan median FA value in the White Matter (WM).

**Figure A.3.**
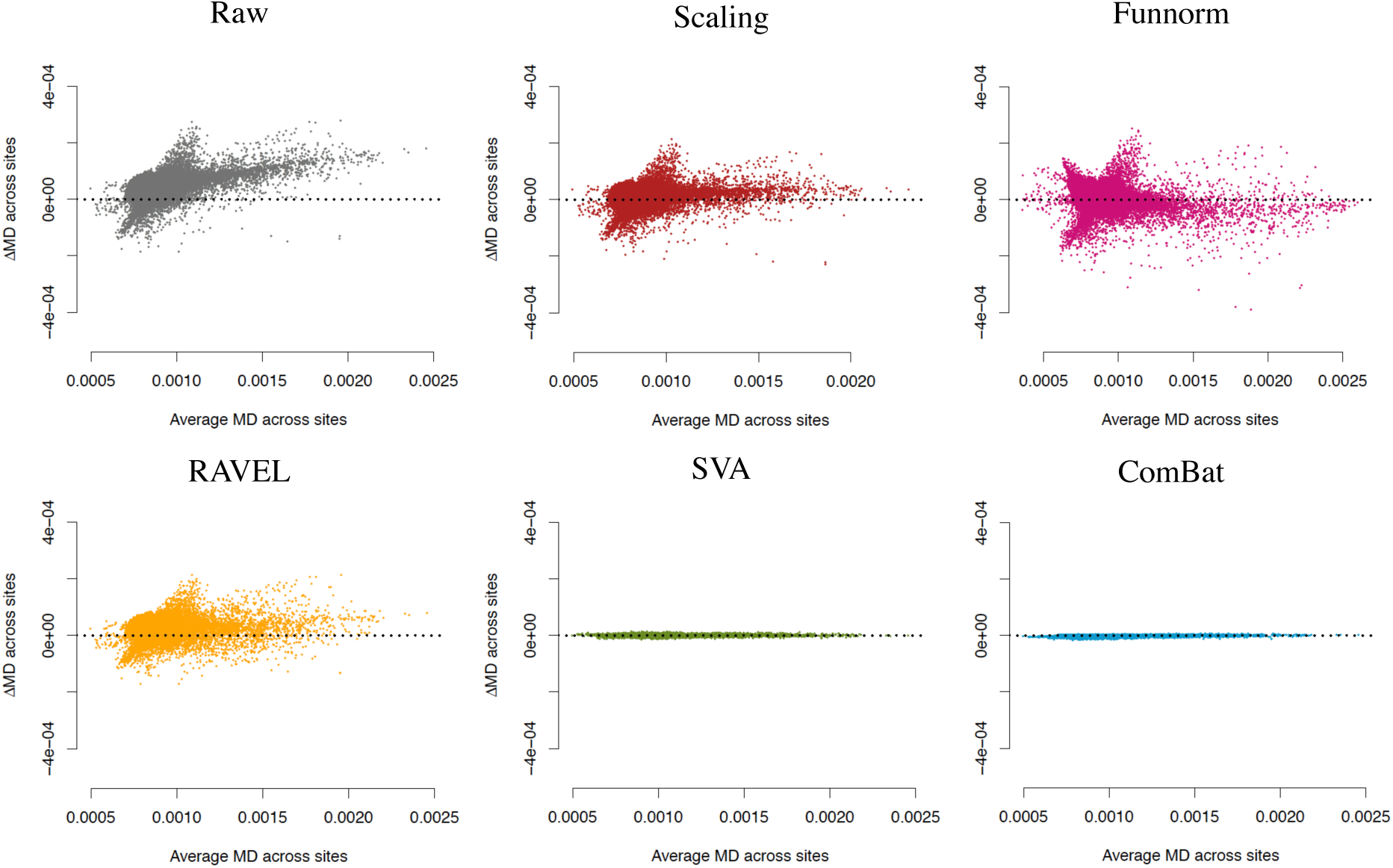
MA-plots for site differences in MD maps. Same as Figure 5, but for MD maps.

**Figure A.4.**
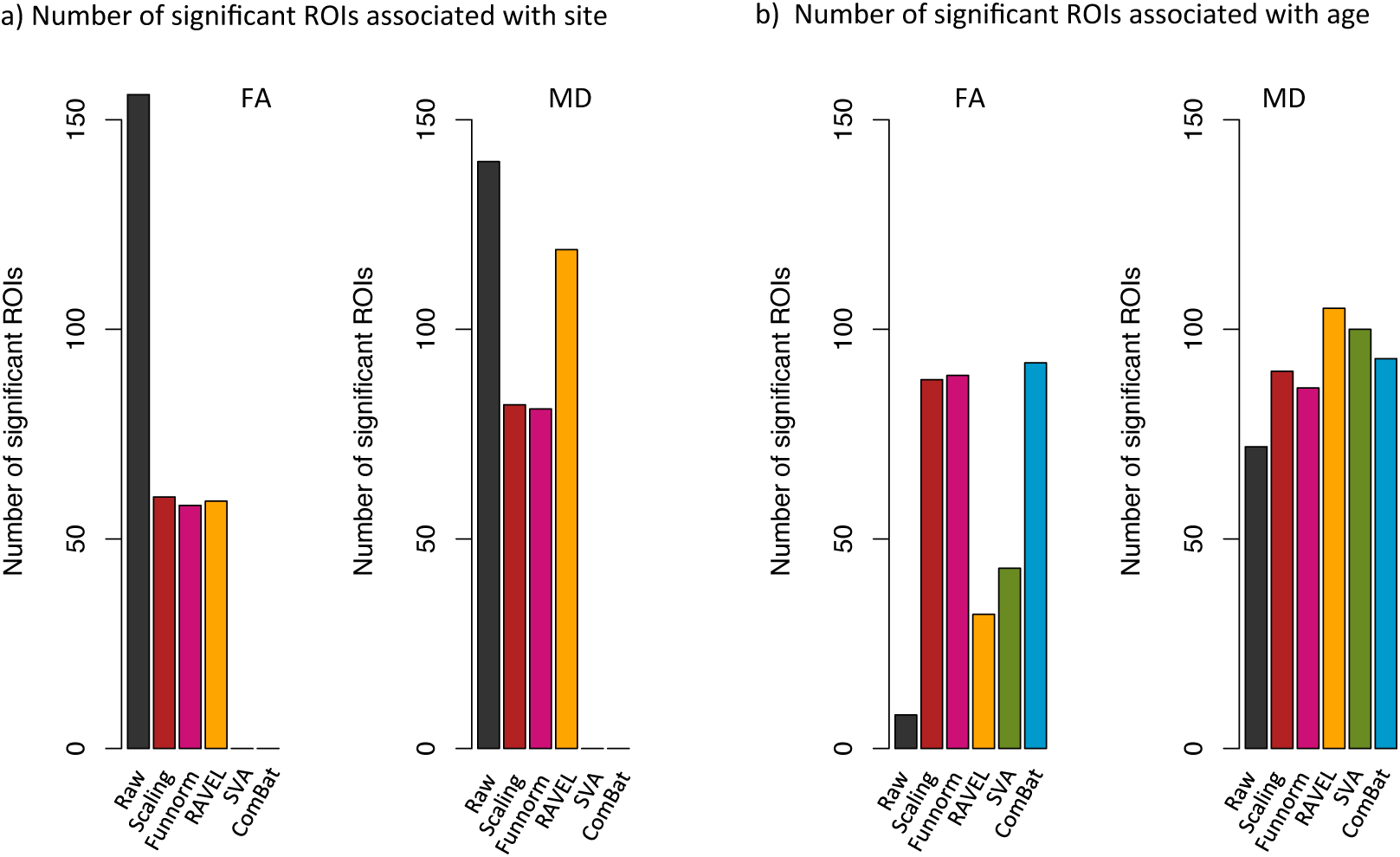
Number of ROIs associated with site and age. Same as Figure 6, but for the 156 regions of interest (ROIs). All p-values were adjusted for multiple comparisons in a conservative manner using Bonferroni correction. (a) In the absence of harmonization (raw data), all 156 ROIs are associated with site in the FA maps, and 140 ROIs are associated with site in the MD maps. Both SVA and ComBat result in 0 ROI associated with site. (b) ComBat performs well at increasing the number of ROIs associated with age (92 ROIs for FA and 92 ROIs for MD), as opposed to 8 ROIs and 72 ROIs in the raw data, for the FA and MD maps respectively.

**Figure A.5.**
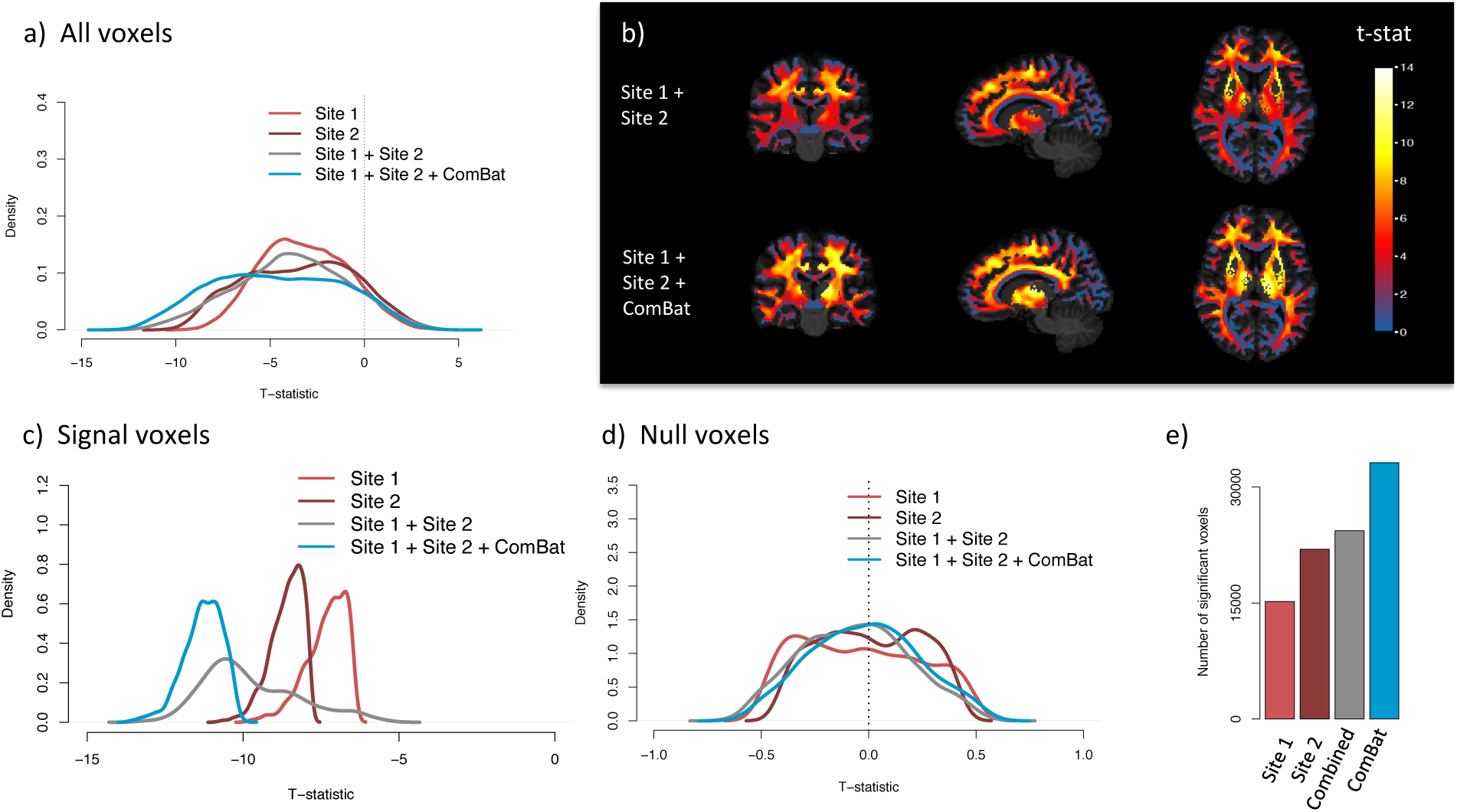
Effect of ComBat harmonization on t-statistics (MD maps). Same as Figure 11, but for the MD maps.

**Figure A.6.**
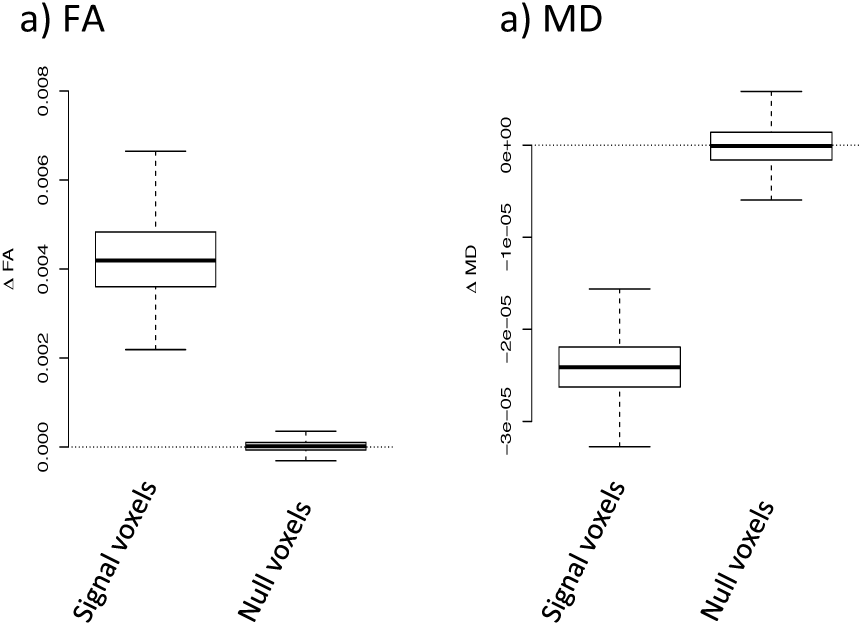
Distribution of the effect sizes for the silver-standards.

**Figure A.7.**
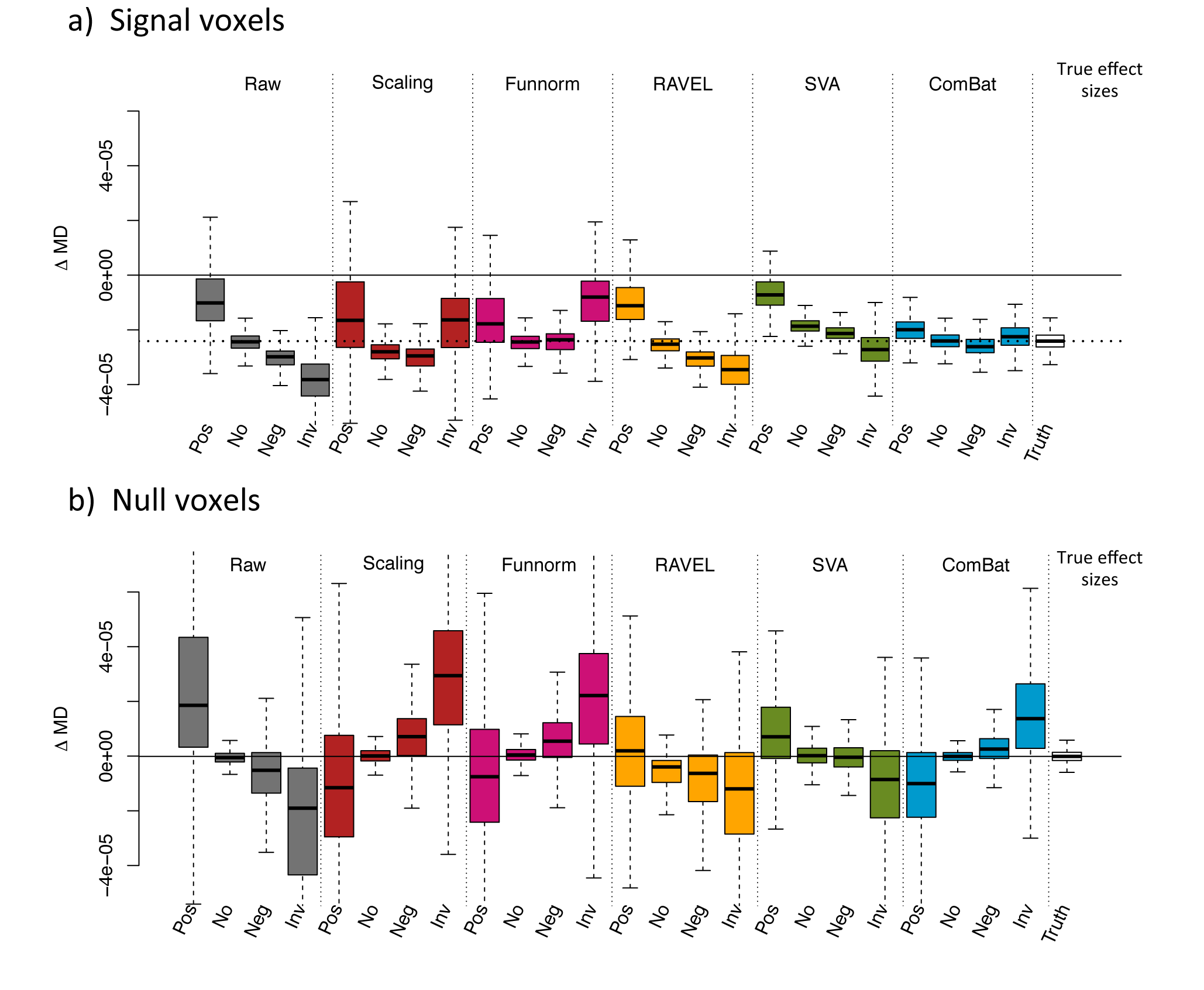
Estimated effect sizes 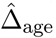 for different confounding scenarios. Same as Figure 10, but for MD.

## References

Andrew L Alexander, Khader M Hasan, Mariana Lazar, Jay S Tsuruda, and Dennis L Parker. Analysis of partial volume effects in diffusion-tensor mri. Magnetic Resonance in Medicine, 45 (5):770–780, 2001.

Andrew L Alexander, Jee Eun Lee, Mariana Lazar, and Aaron S Field. Diffusion tensor imaging of the brain. Neurotherapeutics, 4(3):316–329, 2007.

Manzar Ashtari, Kelly L Cervellione, Khader M Hasan, Jinghui Wu, Carolyn McIlree, Hana Kester, Babak A Ardekani, David Roofeh, Philip R Szeszko, and Sanjiv Kumra. White matter development during late adolescence in healthy males: a cross-sectional diffusion tensor imaging study. Neuroimage, 35(2):501–510, 2007.

Naama Barnea-Goraly, Vinod Menon, Mark Eckert, Leanne Tamm, Roland Bammer, Asya Karchemskiy, Christopher C Dant, and Allan L Reiss. White matter development during childhood and adolescence: a cross-sectional diffusion tensor imaging study. Cerebral cortex, 15(12): 1848–1854, 2005.

Sunita Bava, Rachel Thayer, Joanna Jacobus, Megan Ward, Terry L Jernigan, and Susan F Tapert. Longitudinal characterization of white matter maturation during adolescence. Brain research, 1327:38–46, 2010.

J Martin Bland and DouglasG Altman. Statistical methods for assessing agreement between two methods of clinical measurement. The lancet, 327(8476):307–310, 1986.

B M Bolstad, R A Irizarry, M Astrand, and T P Speed. A comparison of normalization methods for high density oligonucleotide array data based on variance and bias. Bioinformatics, 19(2): 185–193, 2003.

W Cleveland. Visualizing data. at & t bell laboratories, murray hill nj, 1993.

William S. Cleveland. Robust locally weighted regression and smoothing scatterplots. Journal of the American Statistical Association, 74:829–836, 1979.

William S Cleveland. Lowess: A program for smoothing scatterplots by robust locally weighted regression. The American Statistician, 35(1):54, 1981.

Marta Morgado Correia, Thomas A Carpenter, and Guy B Williams. Looking for the optimal dti acquisition scheme given a maximum scan time: are more b-values a waste of time? Magnetic resonance imaging, 27(2):163–175, 2009.

Sergi G Costafreda. Pooling fmri data: meta-analysis, mega-analysis and multi-center studies. Frontiers in neuroinformatics, 3:33, 2009.

Stella J De Wit, Pino Alonso, Lizanne Schweren, David Mataix-Cols, Christine Lochner, José M Menchóon, Dan J Stein, Jean-Paul Fouche, Carles Soriano-Mas, Joao R Sato, et al. Multicenter voxel-based morphometry mega-analysis of structural brain scans in obsessive-compulsive disorder. American journal of psychiatry, 171(3):340–349, 2014.

Sandrine Dudoit, Yee Hwa Yang, Matthew J Callow, and Terence P Speed. Statistical methods for identifying differentially expressed genes in replicated cdna microarray experiments. Statistica sinica, pages 111–139, 2002.

Jean-Philippe Fortin, Aurelie Labbe, Mathieu Lemire, Brent Zanke, Thomas Hudson, Elana Fertig, Celia Greenwood, and Kasper D Hansen. Functional normalization of 450k methylation array data improves replication in large cancer studies. Genome Biology, 15(11):503, 2014. doi:10.1186/s13059-014-0503-2.

Jean-Philippe Fortin, Elizabeth M Sweeney, John Muschelli, Ciprian M Crainiceanu, Russell T Shinohara, Alzheimer’s Disease Neuroimaging Initiative, et al. Removing inter-subject technical variability in magnetic resonance imaging studies. NeuroImage, 132:198–212, 2016a.

Jean-Philippe Fortin, Timothy Triche, and Kasper Hansen. Preprocessing, normalization and integration of the illumina humanmethylationepic array. bioRxiv, 2016b.

J A Gagnon-Bartsch and T P Speed. Using control genes to correct for unwanted variation in microarray data. Biostatistics, 13(3):539–552, 2012. doi:10.1093/biostatistics/kxr034.

Eleftherios Garyfallidis, Matthew Brett, Bagrat Amirbekian, Ariel Rokem, Stefan Van Der Walt, Maxime Descoteaux, and Ian Nimmo-Smith. Dipy, a library for the analysis of diffusion mri data. Frontiers in neuroinformatics, 8:8, 2014.

Yasser Ghanbari, Alex R Smith, Robert T Schultz, and Ragini Verma. Identifying group discriminative and age regressive sub-networks from dti-based connectivity via a unified framework of non-negative matrix factorization and graph embedding. Medical image analysis, 18(8):1337–1348, 2014.

Marco Giannelli, Mirco Cosottini, Maria Chiara Michelassi, Guido Lazzarotti, Gina Belmonte, Carlo Bartolozzi, and Mauro Lazzeri. Dependence of brain dti maps of fractional anisotropy and mean diffusivity on the number of diffusion weighting directions. Journal of Applied Clinical Medical Physics, 11(1), 2009.

Antonio Giorgio, KE Watkins, Martin Chadwick, S James, Louise Winmill, Gwenaëlle Douaud, Nicola De Stefano, Paul M Matthews, Steve M Smith, Heidi Johansen-Berg, et al. Longitudinal changes in grey and white matter during adolescence. Neuroimage, 49(1):94–103, 2010.

Rafael A Irizarry, Daniel Warren, Forrest Spencer, Irene F Kim, Shyam Biswal, Bryan C Frank, Edward Gabrielson, Joe G N Garcia, Joel Geoghegan, Gregory Germino, Constance Griffin, Sara C Hilmer, Eric Hoffman, Anne E Jedlicka, Ernest Kawasaki, Francisco Mart´inez-Murillo, Laura Morsberger, Hannah Lee, David Petersen, John Quackenbush, Alan Scott, Michael Wilson, Yanqin Yang, Shui Qing Ye, and Wayne Yu. Multiple-laboratory comparison of microarray platforms. Nature Methods, 2(5):345–50, 2005. doi:10.1038/nmeth756.

Neda Jahanshad, Peter V Kochunov, Emma Sprooten, Ren´e C Mandl, Thomas E Nichols, Laura Almasy, John Blangero, Rachel M Brouwer, Joanne E Curran, Greig I de Zubicaray, et al. Multi-site genetic analysis of diffusion images and voxelwise heritability analysis: A pilot project of the enigma–dti working group. Neuroimage, 81:455–469, 2013.

Mark Jenkinson and Stephen Smith. A global optimisation method for robust affine registration of brain images. Medical image analysis, 5(2):143–156, 2001.

Mark Jenkinson, Peter Bannister, Michael Brady, and Stephen Smith. Improved optimization for the robust and accurate linear registration and motion correction of brain images. Neuroimage, 17(2):825–841, 2002.

W Evan Johnson, Cheng Li, and Ariel Rabinovic. Adjusting batch effects in microarray expression data using empirical bayes methods. Biostatistics, 8(1):118–127, 2007.

Derek K Jones. The effect of gradient sampling schemes on measures derived from diffusion tensor mri: a monte carlo study. Magnetic Resonance in Medicine, 51(4):807–815, 2004.

Mina Kim, Itamar Ronen, Kamil Ugurbil, and Dae-Shik Kim. Spatial resolution dependence of dti tractography in human occipito-callosal region. Neuroimage, 32(3):1243–1249, 2006.

Kathleen Oros Klein, Stepan Grinek, Sasha Bernatsky, Luigi Bouchard, Antonio Ciampi, Ines Colmegna, Jean-Philippe Fortin, Long Gao, Marie-France Hivert, Marie Hudson, et al. funtoonorm: an r package for normalization of dna methylation data when there are multiple cell or tissue types. Bioinformatics, page btv615, 2015.

Peter Kochunov, Neda Jahanshad, Emma Sprooten, Thomas E Nichols, Ren´e C Mandl, Laura Almasy, Tom Booth, Rachel M Brouwer, Joanne E Curran, Greig I de Zubicaray, et al. Multi-site study of additive genetic effects on fractional anisotropy of cerebral white matter: comparing meta and megaanalytical approaches for data pooling. NeuroImage, 95:136–150, 2014.

Stine K Krogsrud, Anders M Fjell, Christian K Tamnes, Håkon Grydeland, Lia Mork, Paulina Due-Tønnessen, Atle Bjørnerud, Cassandra Sampaio-Baptista, Jesper Andersson, Heidi Johansen-Berg, et al. Changes in white matter microstructure in the developing braina longitudinal diffusion tensor imaging study of children from 4 to 11years of age. NeuroImage, 124:473–486, 2016.

Catherine Lebel and Christian Beaulieu. Longitudinal development of human brain wiring continues from childhood into adulthood. The Journal of Neuroscience, 31(30):10937–10947, 2011.

Jeffrey T Leek and Roger D Peng. Opinion: Reproducible research can still be wrong: adopting a prevention approach. Proc Natl Acad Sci U S A, 112(6):1645–6, Feb 2015. doi:10.1073/pnas.1421412111.

Jeffrey T Leek and John D Storey. Capturing heterogeneity in gene expression studies by surrogate variable analysis. PLoS Genetics, 3(9):1724–1735, 2007. doi:10.1371/journal.pgen.0030161.

Jeffrey T Leek and John D Storey. A general framework for multiple testing dependence. Proceedings of the National Academy of Sciences, 105(48):18718–18723, 2008. doi:10.1073/pnas.0808709105.

Jeffrey T Leek, W Evan Johnson, Hilary S Parker, Andrew E Jaffe, and John D Storey. The sva package for removing batch effects and other unwanted variation in high-throughput experiments. Bioinformatics, 28(6):882–883, 2012. doi:10.1093/bioinformatics/bts034.

Kristin A Linn, Bilwaj Gaonkar, Jimit Doshi, Christos Davatzikos, and Russell T Shinohara. Addressing confounding in predictive models with an application to neuroimaging. The international journal of biostatistics, 12(1):31–44, 2016a.

Kristin A Linn, Bilwaj Gaonkar, Theodore D Satterthwaite, Jimit Doshi, Christos Davatzikos, and Russell T Shinohara. Control-group feature normalization for multivariate pattern analysis of structural mri data using the support vector machine. NeuroImage, 132:157–166, 2016b.

H Mirzaalian, L Ning, P Savadjiev, O Pasternak, S Bouix, O Michailovich, G Grant, CE Marx, RA Morey, LA Flashman, et al. Inter-site and inter-scanner diffusion mri data harmonization. NeuroImage, 135:311–323, 2016.

John Muschelli, Elizabeth M Sweeney, Martin A Lindquist, and Ciprian M Crainiceanu. fslr: Connecting the fsl software with r. The R Journal, 7(1):163–175, Feb 2015.

Yangming Ou, Aristeidis Sotiras, Nikos Paragios, and Christos Davatzikos. Dramms: Deformable registration via attribute matching and mutual-saliency weighting. Medical image analysis, 15 (4):622–639, 2011.

Giovanni Parmigiani, Elizabeth S Garrett, Rafael A Irizarry, and Scott L Zeger. The analysis of gene expression data: an overview of methods and software. In The analysis of gene expression data, pages 1–45. Springer, 2003.

Gholamreza Salimi-Khorshidi, Stephen M Smith, John R Keltner, Tor D Wager, and Thomas E Nichols. Meta-analysis of neuroimaging data: a comparison of image-based and coordinate-based pooling of studies. Neuroimage, 45(3):810–823, 2009.

Theodore D Satterthwaite, Mark A Elliott, Kosha Ruparel, James Loughead, Karthik Prabhakaran, Monica E Calkins, Ryan Hopson, Chad Jackson, Jack Keefe, Marisa Riley, et al. Neuroimaging of the philadelphia neurodevelopmental cohort. Neuroimage, 86:544–553, 2014.

Vincent J Schmithorst, Marko Wilke, Bernard J Dardzinski, and Scott K Holland. Correlation of white matter diffusivity and anisotropy with age during childhood and adolescence: A cross-sectional diffusion-tensor mr imaging study 1. Radiology, 222(1):212–218, 2002.

Russell T Shinohara, Elizabeth M Sweeney, Jeff Goldsmith, Navid Shiee, Farrah J Mateen, Peter A Calabresi, Samson Jarso, Dzung L Pham, Daniel S Reich, Ciprian M Crainiceanu, Australian Imaging Biomarkers Lifestyle Flagship Study of Ageing, and Alzheimer’s Disease Neuroimaging Initiative. Statistical normalization techniques for magnetic resonance imaging. Neuroimage Clin, 6:9–19, 2014. doi:10.1016/j.nicl.2014.08.008.

Stephen M Smith. Fast robust automated brain extraction. Hum Brain Mapp, 17(3):143–55, Nov 2002. doi:10.1002/hbm.10062.

Christian K Tamnes, Ylva Østby, Anders M Fjell, Lars T Westlye, Paulina Due-Tønnessen, and Kristine B Walhovd. Brain maturation in adolescence and young adulthood: regional age-related changes in cortical thickness and white matter volume and microstructure. Cerebral cortex, 20 (3):534–548, 2010.

Antonio Tristán-Vega and Santiago Aja-Fernández. Dwi filtering using joint information for dti and hardi. Medical image analysis, 14(2):205–218, 2010.

Jessica A Turner. The rise of large-scale imaging studies in psychiatry. GigaScience, 3(1):29, 2014.

Neeltje EM van Haren, Wiepke Cahn, Hilleke E Hulshoff Pol, Hugo G Schnack, Esther Caspers, Adriaan Lemstra, Margriet M Sitskoorn, Durk Wiersma, Rob J van den Bosch, Peter M Dinge-mans, et al. Brain volumes as predictor of outcome in recent-onset schizophrenia: a multi-center mri study. Schizophrenia Research, 64(1):41–52, 2003.

Liang Zhan, Alex D Leow, Neda Jahanshad, Ming-Chang Chiang, Marina Barysheva, Agatha D Lee, Arthur W Toga, Katie L McMahon, Greig I de Zubicaray, Margaret J Wright, et al. How does angular resolution affect diffusion imaging measures? Neuroimage, 49(2):1357–1371, 2010.

Y Zhang, M Brady, and S Smith. Segmentation of brain mr images through a hidden markov random field model and the expectation-maximization algorithm. IEEE Trans Med Imaging, 20 (1):45–57, Jan 2001. doi:10.1109/42.906424.

Tong Zhu, Xiaoxu Liu, Michelle D Gaugh, Patrick R Connelly, Hongyan Ni, Sven Ekholm, Giovanni Schifitto, and Jianhui Zhong. Evaluation of measurement uncertainties in human diffusion tensor imaging (dti)-derived parameters and optimization of clinical dti protocols with a wild bootstrap analysis. Journal of Magnetic Resonance Imaging, 29(2):422–435, 2009.

Tong Zhu, Rui Hu, Xing Qiu, Michael Taylor, Yuen Tso, Constantin Yiannoutsos, Bradford Navia, Susumu Mori, Sven Ekholm, Giovanni Schifitto, et al. Quantification of accuracy and precision of multi-center dti measurements: a diffusion phantom and human brain study. Neuroimage, 56 (3):1398–1411, 2011.

